# Deciphering Endothelial and Mesenchymal Organ Specification in Vascularized Lung and Intestinal Organoids

**DOI:** 10.1101/2024.02.06.577460

**Authors:** Yifei Miao, Cheng Tan, Nicole M. Pek, Zhiyun Yu, Kentaro Iwasawa, Daniel O. Kechele, Nambirajan Sundaram, Victor Pastrana-Gomez, Keishi Kishimoto, Min-Chi Yang, Cheng Jiang, Jason Tchieu, Jeffrey A. Whitsett, Kyle W. McCracken, Robbert J. Rottier, Darrell N. Kotton, Michael A. Helmrath, James M. Wells, Takanori Takebe, Aaron M. Zorn, Ya-Wen Chen, Minzhe Guo, Mingxia Gu

## Abstract

To investigate the co-development of vasculature, mesenchyme, and epithelium crucial for organogenesis and the acquisition of organ-specific characteristics, we constructed a human pluripotent stem cell-derived organoid system comprising lung or intestinal epithelium surrounded by organotypic mesenchyme and vasculature. We demonstrated the pivotal role of co-differentiating mesoderm and endoderm via precise BMP regulation in generating multilineage organoids and gut tube patterning. Single-cell RNA-seq analysis revealed organ specificity in endothelium and mesenchyme, and uncovered key ligands driving endothelial specification in the lung (e.g., WNT2B and Semaphorins) or intestine (e.g., GDF15). Upon transplantation under the kidney capsule in mice, these organoids further matured and developed perfusable human-specific sub-epithelial capillaries. Additionally, our model recapitulated the abnormal endothelial-epithelial crosstalk in patients with *FOXF1* deletion or mutations. Multilineage organoids provide a unique platform to study developmental cues guiding endothelial and mesenchymal cell fate determination, and investigate intricate cell-cell communications in human organogenesis and disease.

**Highlights:**

- BMP signaling fine-tunes the co-differentiation of mesoderm and endoderm.
- The cellular composition in multilineage organoids resembles that of human fetal organs.
- Mesenchyme and endothelium co-developed within the organoids adopt organ-specific characteristics.
- Multilineage organoids recapitulate abnormal endothelial-epithelial crosstalk in FOXF1-associated disorders.

## Introduction

The proper development of organs and maintenance of tissue homeostasis rely heavily on the precise establishment of the vascular bed and mesenchymal structure^1–3^. Recent studies have created single-cell atlases of human fetal^4^ and adult organs^5^, revealing unique organotypic features in the capillary endothelium and surrounding mesenchyme. It is now evident that the vasculature serves not only as support and nutrient transport systems but also wields significant influence in orchestrating organ formation, maturation, and post-damage regeneration through intricate signal communication pathways. To investigate the spatial-temporal dynamics of this biological process, it is critical to develop *in vitro* platforms that incorporate organotypic vasculature and relevant cell lineages with appropriate cellular architecture.

During embryonic development, the definitive endoderm (DE), which is differentiated from the pluripotent epiblast, gives rise to the vast majority of highly specialized epithelial cell types lining the respiratory and digestive gut tube systems^6–8^. Mesodermal lineages arising from the same epiblast form mesenchymal tissues and vasculature, which inductively pattern the gut tube organs^6,9^. To generate complex organoids, assembly strategies have been developed through fusion of multiple cell types subsequent to their separate differentiation^10,11^. Nevertheless, this strategy lacks the initial interactions among multiple germ layers, which is critical for the proper differentiation of progenitor populations that ultimately give rise to diverse cell types within an organ. In our study, utilizing a precisely controlled mesoderm-endoderm co-differentiation approach regulated by BMP signaling, we generated multicellular gut tube organoids derived from human pluripotent stem cells (hPSCs), including DE-derived lung or intestine epithelium and lateral plate mesoderm (LPM)-derived mesenchyme and vasculature. Notably, the vascularized organoids facilitated the emergence of organ-specific endothelium and mesenchyme. This not only allows for the investigation of the environmental cues driving mesodermal cell fate determination, but also provides a complex platform to study the impact of vascular and mesenchymal abnormalities on surrounding cell types in various congenital disorders.

## Results

### Fine-tuning mesoderm-endoderm co-differentiation by manipulating BMP signaling

To generate vascularized lung and intestinal organoids from hPSCs (Fig. 1A), we recapitulated the process of human early embryonic development by concurrently differentiating the corresponding germ layer progenitors within an intact 3D structure. Within the first three days, hPSCs-derived embryonic bodies (EBs) were differentiated to spheroids consisting of both DE (FOXA2^+^ or CXCR4^+^/KIT^+^) and mesoderm (HAND1^+^) lineages (Fig. 1A and Fig. S1). It has been reported that BMP4 plays a pivotal role in initiating the Wnt-Nodal hierarchy, prompting the self-organization of hESCs to pattern both embryonic and extraembryonic germ layers within a flat, geometrically confined disc^12,13^. In the 3D EB system, we found that Wnt activation by CHIR99021 (CHIR) from day 0-1 was indispensable for priming the co-differentiation process (Fig. S1A-S1D). The balance between DE and mesoderm was primarily influenced by the activation of Nodal through Activin A and Wnt signaling (Fig. 1A and Fig. S1B, S1C, and S1E)^8^. A three-day treatment of Activin A was sufficient to induce high-purity DE^14^, while prolonged activation of the Wnt pathway for over 24 hours significantly diverted DE toward the mesoderm lineage (Fig. 1A and Fig. S1D-S1E). Next, to generate spheroids encompassing both DE and mesoderm, we manipulated additional signaling pathways. Unlike on the 2D platform, FGF2 activation or PI3K inhibition played a negligible role within the EBs in terms of germ layer specification (Fig. S1F-S1G)^8^. Notably, we found that the ratio of DE and mesoderm could be fine-tuned by varying the duration of BMP activation-prolonged BMP4 treatment duration suppressed DE while enhanced HAND1^+^ mesoderm (Fig. 1B and Fig. S1E and S1H).

**Fig. 1.**
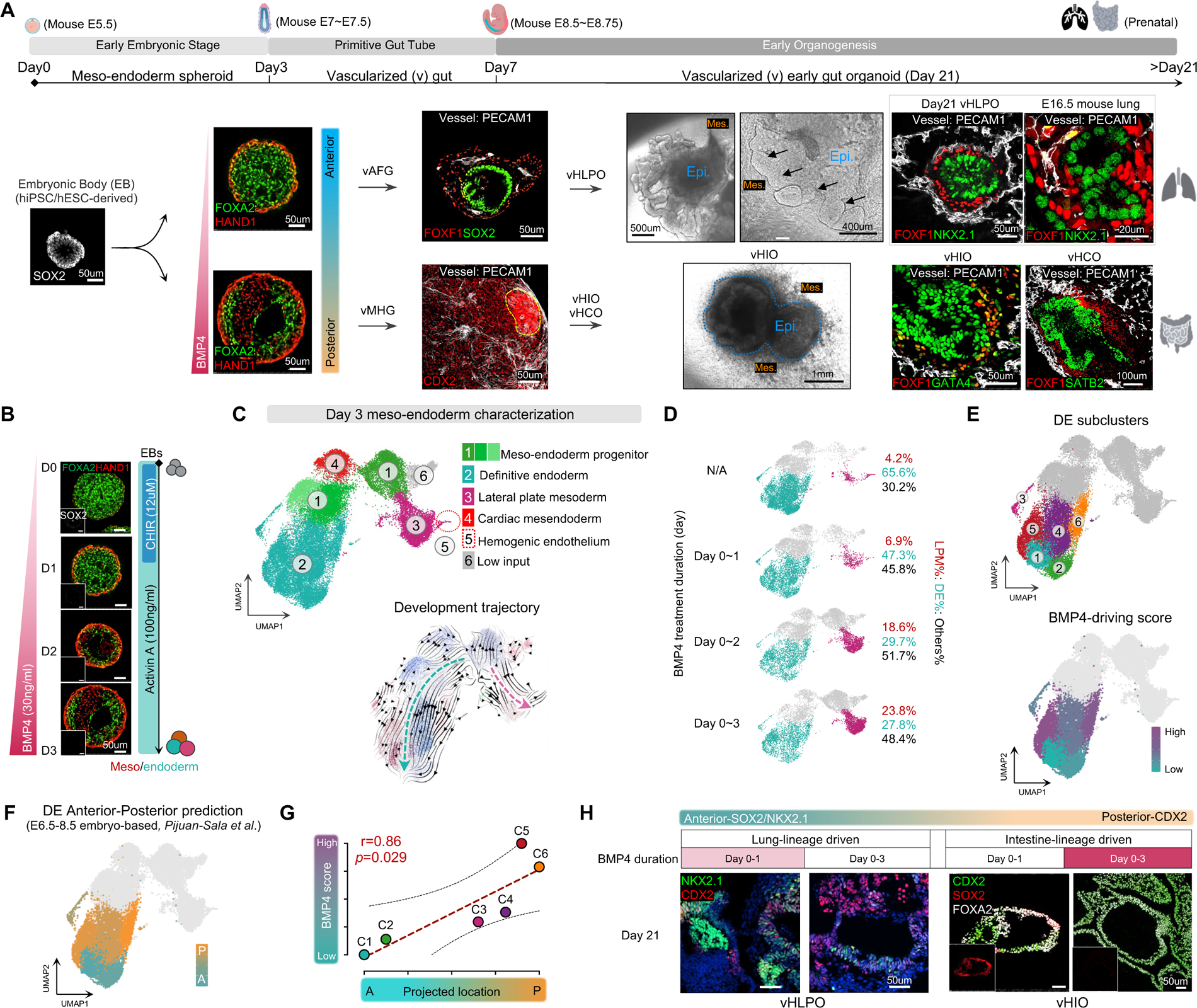
Refining BMP regulation for endoderm-mesoderm co-differentiation, gut tube patterning, and generating vascularized lung and intestinal organoids. A. Schematic illustration and immunostaining of vascularized organoids at the key developmental stages. Regulated by different durations of BMP activation, hPSC-derived EBs were differentiated into meso-endoderm spheroids by day 3, vascularized foregut or midgut/hindgut by day 7, and vascularized lung or intestinal progenitor organoids by day 21. hiPSC, human induced pluripotent stem cell; hESC, human embryonic stem cell; EB: embryonic body; vAFG, vascularized anterior foregut; vMHG: vascularized midgut/hindgut; vHLPO, vascularized human lung progenitor organoid (NKX2.1^+^ Epi.); vHIO, vascularized human intestinal organoid (GATA4^+^ Epi.); vHCO, vascularized human colonic organoid (SATB2^+^ Epi.); Epi., epithelium; Mes., mesenchyme. Yellow dashed circle: MHG epithelium. Black arrows: lung bud-like structure. B. Whole-mount staining of day-3 spheroids subjected to different durations of BMP4 treatment. FOXA2: Definitive endoderm (DE); HAND1: lateral plate mesoderm; SOX2: pluripotent stem cells. C. UMAP projection and RNA velocity analysis of day 3 meso-endoderm spheroids. Black arrows indicated the developmental trajectory directions determined by velocyto algorithm. D. UMAP distributions of DE and LPM under different durations of BMP4 treatment. E. Highlight of Fig. 1C DE subclusters and corresponding BMP-driving score, which is calculated based on the distribution proportions of each sample subjected to different BMP4 treatment durations. F. Projection of anterior and posterior mouse gut subtypes (Pijuan-Sala et al.,) onto the UMAP of human spheroid DE subclusters. G. Correlation between the A-P projection of the day 3 human spheroid DE subclusters and BMP-driving scores. H. Immunostaining conducted on day 21 vHLPO (lung epithelium: NKX2.1+, CDX2-) and vHIO (intestinal epithelium: CDX2+, FOXA2+, SOX2-) showed that the durations of BMP4 exposure within the initial three days of differentiation predetermined the anterior versus posterior epithelial cell fate by day 21. See also Figure S1, S2, and Table S1.

Single-cell RNA-seq (scRNA-seq) on these day-3 meso-endoderm spheroids further demonstrated the simultaneous emergence of DE and mesoderm lineages (Fig. 1C, Fig. S2A-S2D, and Table S1). The pseudotime trajectory analysis suggested that they were developmentally continuous from the heterogenous meso-endoderm progenitors (Fig. 1C). Additionally, cardiac mesendoderm and endothelial progenitors co-emerged (Fig. S2B and S2E) alongside endoderm, reinforcing the cross-lineage communications between the heart and adjacent endoderm. This simultaneous emergence suggests a synchronized coordination in their distinct developmental pathways^15,16^. To benchmark the meso-endoderm spheroids to their *in vivo* counterparts, scRNA-seq data from human embryo at the similar Carnegie Stage (CS) 7^17^ was projected onto day-3 spheroids (Fig. S2F): the meso-endoderm progenitors appeared akin to early emergent mesoderm, while the LPM resembled advanced mesoderm.

scRNA-seq analyses of day-3 spheroids exposed to varying BMP4 treatments spanning from 0 to 3 days further supported the precise controlling of DE versus mesodermal population ratio via fine-tuning BMP durations (Fig. 1D). The addition of BMP4 significantly induced the LPM percentage from 4.2% to 23.8%, while reducing the DE percentage from 65.6% to 27.8%. The cellular heterogenicity was further amplified in DE subclusters, which was primarily determined by BMP signaling (Fig. 1E and Fig. S2G). More interestingly, we found these diverse DE subclusters exerted distinct anterior-posterior patterns as demonstrated by fractional projection to E6.5∼E8.5 mouse primitive gut regional features^18^ (Fig. 1F-G and Fig. S2H-S2K) and the expression of known anterior and posterior endoderm markers^7^ (Fig. S2L-S2M). Interestingly, this imprinted regional signature driven by BMP during the early embryonic stage predisposed the anterior-posterior early gut tube pattern in the subsequent organogenesis phase (day 7-21) (Fig. 1H): short activation of BMP signaling from day 0-1 facilitated the subsequent lung lineage specification at day 21, whereas prolonged BMP stimulation from day 0-3 resulted in a posterior intestinal fate.

Collectively, by adjusting the duration of BMP activation during the first three days of differentiation, we established a 3D differentiation system to co-develop DE and mesoderm populations, providing all essential progenitors for generating epithelial, endothelial, and mesenchymal cells within the same spheroid during subsequent stages.

### Generation of vascularized lung and intestinal organoids

After the early embryogenic stage, a series of morphogens transform the DE into a primitive gut tube surrounded by mesoderm, which is regionalized into the foregut, midgut, and hindgut domains along the anterior-posterior (A-P) axes^7^. To generate vascularized anterior foregut (vAFG) and vascularized midgut/hindgut (vMHG) organoids from meso-endoderm spheroids from day 3 to day 7, we first examined previously reported key signaling for epithelial differentiation without mesoderm induction (Fig. 1A, Fig. 2A-2E, and Fig. S3A-S3F). Inhibition of TGFb by SB-431542 had minimal impact on either AFG purity or mesenchymal (FOXF1^+^)/vessel (PECAM1^+^) induction (Fig. 2A). Additionally, we examined the role of hedgehog (HH) activation in our system, as it has been shown that HH activation promoted the expansion of lung (AFG) endoderm and mesenchyme^19^. Although we observed that HH activation by SAG resulted in an increase in mesenchymal population in day-7 vAFG (Fig. 2A), it significantly dampened the differentiation of vAFG into NKX2.1^+^ lung epithelium in day 21 vascularized human lung progenitor organoids (vHLPO) (Fig. 2C). This data suggested that endogenous HH signaling within the multilineage organoid was sufficient for lung organoid differentiation, whereas exogenous HH may compromise the purity of lung epithelial differentiation. It has been shown that Wnt and FGF activation combined with BMP and TGFb inhibition led to the formation of FG organoids, which will be further differentiated into lung and gastric organoids^19–21^. However, we found that in our meso-endodermal co-differentiation protocol, activating Wnt (CHIR) and FGF (FGF4) along with BMP inhibition (Noggin) resulted in the posterization (CDX2^+^) of the AFG (SOX2^+^) (Fig. 2B). Collectively, due to the sufficient expression of endogenous morphogens in 3D multilineage spheroids, our protocol is notably simpler compared to prior methods^22–24^, requiring only BMP inhibition for properly patterning the vAFG (days 3-7) to specify the lung lineage (days 7-21) (Fig. S4).

**Fig. 2.**
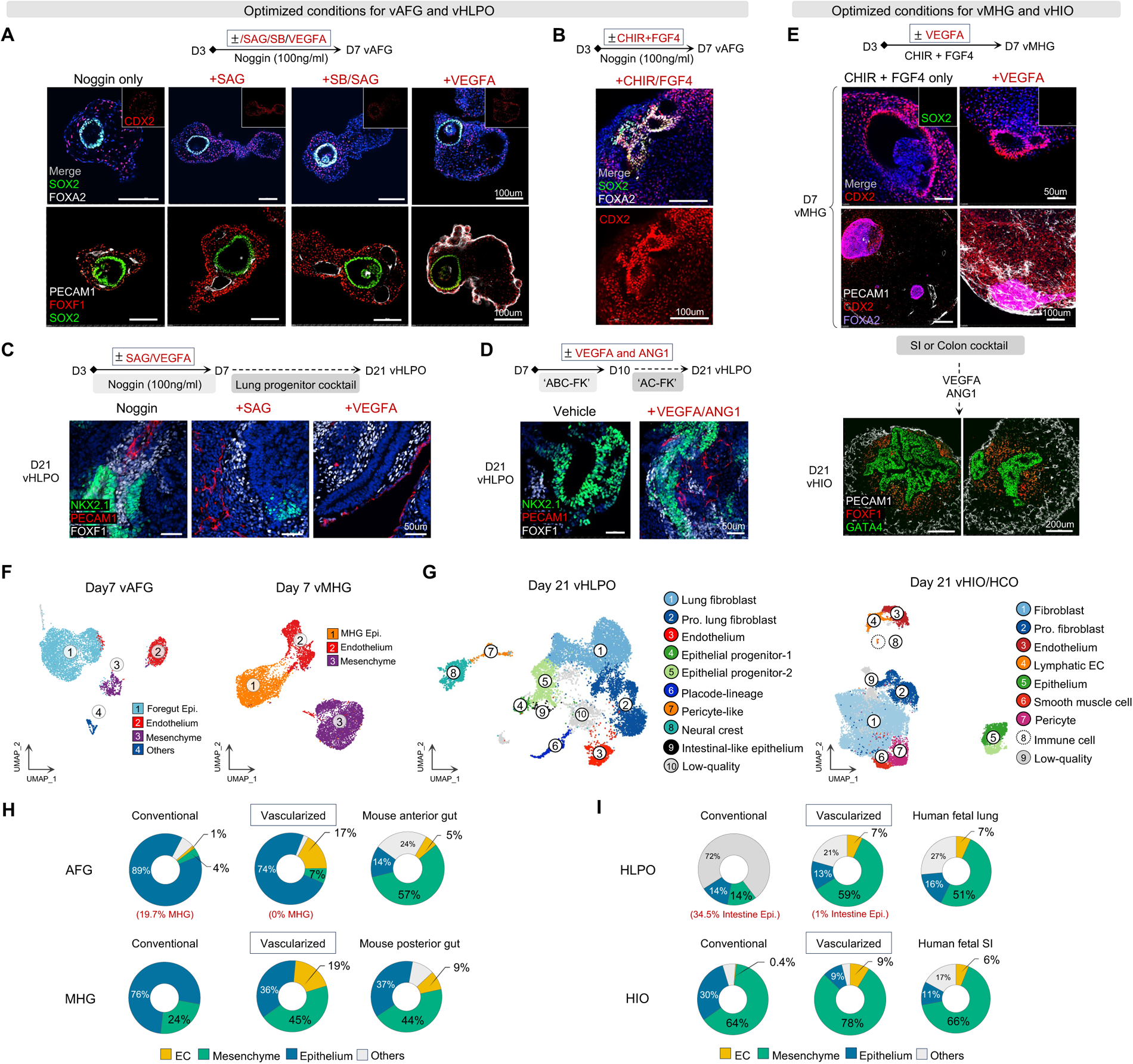
Generation and characterization of the vascularized organoids. **A.** Spheroids were treated with SAG (1uM), SB431542 (10uM), or VEGFA (100ng/ml) from day 3 to day 7, followed by immunostaining on day 7 for anterior gut (SOX2/FOXA2), posterior gut (CDX2/FOXA2), endothelial (PECAM1), and mesenchymal (FOXF1) markers. **B.** Spheroids were treated with CHIR (2uM)+FGF4 (500ng/ml) from day 3 to day 7, followed by immunostaining on day 7 for anterior gut (SOX2/FOXA2) and posterior gut (CDX2/FOXA2) markers. **C**. Spheroids were treated with SAG (1uM) or VEGFA (100ng/ml) from day 3 to day 7, followed by immunostaining on day 21 for lung epithelial (NKX2.1), endothelial (PECAM1), and mesenchymal (FOXF1) markers. **D.** Spheroids were exposed to VEGFA (100ng/ml) and ANG1 (100ng/ml) from day 7 to day 21, followed by immunostaining on day 21 for lineage-specific makers. **E.** Spheroids from day 3 to day 7 were tested for VEGFA (100ng/ml), and vessels were further maintained via supplementing VEGFA (100ng/ml) and ANG1 (100ng/ml) from day 7 to day 21. Immunostaining on day 7 and day 21 organoids for epithelial (CDX2, FOXA2, GATA4), endothelial (PECAM1), and mesenchymal (FOXF1) markers. **F.** UMAP projection of day-7 vAFG and vMHG. Epi.: epithelium. **G.** UMAP projection of day-21 vHLPO and vHIO/vHCO. Pro.: proliferation. **H.** Cell type distributions in day-7 vAFG and vMHG, conventional AFG (Hein et al.) and MHG (Holloway et al.), and E9.0 mouse embryonic anterior (Han et al.) and E8.75 posterior gut (Nowotschin et al.). **I.** Cell type distributions in day-21 vHLPO and vHIO/vHCO, conventional human lung bud organoid (Hein et al.) and human intestinal organoid (Holloway et al.), and human fetal lung (PCW 11 week, He et al.) and small intestine (PCW 10 week, Yu et al.). See also Figure S3, S4.

Next, we investigated the effectiveness and timing of supplementing angiogenic factors to induce vascular formation within gut tube organoids. We found that introducing VEGFA during the primitive gut tube phase (day 3-7) expanded the endogenous pool of ECs without affecting AFG epithelial purity and quantity (Fig. 2A and Fig. S3A). However, this intervention led to the suppression of lung epithelial generation (NKX2-1^+^) at a later stage (Fig. 2C). In contrast, the combined application of VEGFA and angiopoietin 1 (ANG1) synergistically augmented endothelial differentiation during the organogenesis phase (day 7∼21), while maintaining the purity of lung epithelial differentiation (Fig. 2D). ANG1 was only introduced at a later stage due to its significant role in fostering the organization and maturation of newly formed vessels and maintaining the structural integrity of adult vasculature^25^.

To generate vHLPO, conventional lung organoid progenitor cocktail^22,26^ (three days of all-trans retinoic acid (ATRA)/BMP4/CHIR/FGF7/FGF10-‘ABC-FK’, followed by ‘AC-FK’ till day 21.) was added in combination with VEGFA and ANG1 (Fig. 2D). This method patterned vAFG into vHLPO containing spatially organized lung epithelium (NKX2.1^+^), mesenchyme (FOXF1^+^), and ECs (PECAM1^+^), which were morphologically reminiscent to E16.5 mouse lung structure (Fig. 1A).

To generate vascularized human intestinal organoids (vHIO) and colonic organoids (vHCO), day three meso-endodermal spheroids were exposed to CHIR and FGF4 from day 3 to day 7^27,28^. This process yielded pure CDX2^+^ posterior MHG while concomitantly differentiating mesenchyme and ECs (Fig. 2E). Starting from day 3, the combination of high-dose VEGFA with conventional intestinal epithelial differentiation cocktails^29^ increased EC generation in day-7 vMHG (Fig. 2E and Fig. S3B-S3C) and day-21 vHIOs (Fig. 2E) without compromising posterior gut identity (Fig. S3E). High dose of VEGFA and ANG1 were introduced from day7 to 21 to maintain and further mature the vasculature (Fig. 2E). Additionally, the similar regimen could be combined with colonic organoid differentiation protocol^27^ to produce vHCO (Fig. S3F) and there was no adverse impact of introducing angiogenic factors on posterior SI or colonic gut lineage identity (Fig. S3F).

In contrast to prior methods primarily focusing on producing lung, intestine, and colon epithelial lineages, our approach effectively controlled the concurrent development of defined DE and mesodermal lineages within the same spheroid, resulting in the formation of multicellular organoid structures abundant in vasculature with well-organized cellular arrangement (Fig. S4).

### Enhanced cellular heterogeneity in vascularized organoids

To further reveal the cellular heterogeneity of the vascularized organoids, we performed scRNA-seq on day-7 vAFG and vMHG organoids. We focused on the vAFG-‘B1’ and vMHG-‘B3’ groups, given their optimal induction of epithelial, endothelial, and mesenchymal populations on day 7 and day 21 organoids (Fig. 1H and Fig. S3G-S3H). Both day-7 vAFG and vMHG showed expected anterior foregut or posterior midgut/hindgut epithelial features (Fig. S3G-S3H), with appreciable amounts of vascular and mesenchymal populations (Fig. 2F and Fig. S3G-S3H).

Next, we analyzed scRNA-seq data from day 21 vHLPO, vHIO, and vHCO derived from day-7 vAFG and vMHG organoids (Fig. 2G and Fig. S3I-S3P). Based on the human fetal lung single-cell atlas^30^, heterogeneous epithelial, mesenchymal, and endothelial populations were identified within vHLPO (Fig. S3I-S3N). For example, fibroblasts in vHLPO were similar to human early fibroblast, mesenchyme, and myofibroblast (Fig. S3K); cluster 6 fibroblast resembled a subset of human myofibroblast surrounding the stalk epithelium, which was enriched with Wnt-responsive genes^30^ (*LEF1* and *NOTUM,* Fig. S3L). The ECs within vHLPO represented early human fetal lung capillaries (Fig. S3M), while cluster 12 lung epithelial progenitor-1 expressed human fetal lung tip-like markers^30^ (*SOX9* and *TPPP3,* Fig. S3I) and cluster 3 lung epithelial progenitor-2 shared certain mesenchymal features (Fig. S3J).

The scRNA-seq analysis conducted on the vHIOs and vHCOs also unveiled multiple cell types, including epithelium, mesenchyme, and ECs (Fig. 2G and Fig. S3O-S3P). Notably, a distinctive group of lymphatic ECs (LECs, *LYVE1*+/*PROX1*+, Fig. S3O) and a small cluster of immune cells (Fig. S3P) spontaneously co-emerged^31^. These cells provide a unique *in vitro* platform to investigate the development of the lymphatic system, a crucial player in dietary lipid transportation and immunosurveillance to maintain intestinal homeostasis^32^.

By timely introducing proper angiogenic factors, our current protocol increased EC differentiation in day-7 vAFG (17%), day-7 vMHG (19%), day-21 vHLPO (7%), and day-21 vHIOs and vHCOs (9%, Fig. 2H-2I), which were significantly improved as compared with conventional day-10 AFG^33^ (1%), day-8 hindgut^29^ (0%), day-31 HLPO^33^ (0%), and day-22 HIO organoids^29^ (0.4%, Fig. 2H-2I). More importantly, the co-emergence of appropriate vasculature and mesenchyme within vascularized organoids also improved anterior epithelial lineage purity. In vAFG, there was barely any MHG contamination (0%) as compared to the conventional AFG differentiation protocol (19.7%, Fig. 2H). Consequently, there was 34.5% of hindgut epithelium within conventional HLPO, while only 1% of intestinal epithelium was found in vHLPO (Fig. 2I). Next, we explored the cell composition similarity between vascularized organoids and their *in vivo* counterparts. The cell composition distributions within vMHG organoids closely mirrored those found in the E8.75 mouse posterior gut atlas^34^ (Fig. 2H), while vAFG organoids contained less mesenchyme and more EC and epithelium as compared to E9.0 mouse anterior gut^9^ (Fig. 2H). Consequently, by day 21, cell compositions in both vHLPO and vHIO organoids closely resembled those observed in human fetal lung^30^ and small intestine tissue^35^ at early developmental stage PCW 10-11 weeks (Fig. 2I).

Overall, our novel vascularized organoid model not only encompassed essential cell types crucial for early organogenesis but also exhibited a striking resemblance to the cellular compositions found in the embryonic gut tubes of both humans and mice.

### Maturation of vascularized lung and intestinal organoids in vivo

To enhance the maturation and perfusion of the vascularized gut organoids, we transplanted them under the kidney capsule of immunocompromised mice for three months^14^ (Fig. 3A). Prior to transplantation, we further differentiated the organoids *in vitro* until day 39. In addition to VEGFA and ANG1, vHLPO was treated with a cocktail designed for lung branching and distalization, including CHIR, Dexamethasone, cAMP, IBMX, FGF7, and FGF10^22^. Meanwhile, vHIO and vHCO were further matured with EGF. Following transplantation, we observed abundant vascular formation (Fig. 3B) and perfusable subepithelial capillary beds that were filled with red blood cells (Fig. 3C and Fig. S5A-S5B) in transplanted (t) vascularized human lung organoids (tvHLuO), tvHIOs, and tvHCOs. Notably, these structures were rarely presented in the conventional epithelial organoids after transplantation^23^ (Fig. 3D and Fig. S5C-S5D). Interestingly, most of the vasculature adjacent to the organoid epithelium originated from human hPSC rather than the host (Fig. 3D-3E and Fig. S5E). Notably, the presence of perfused vessels based on anastomotic structures served as a hallmark for improved organoid maturation and engraftment *in vivo* (Fig. 3F). Additionally, we observed a greater variety of epithelial cell types in tvHLuO, tvHIO, and tvHCO after transplantation (Fig. S5F-S5G). Within the tvHLuO, we profiled the proteomics from the secreted fluid and discovered all the major lung secretory products, such as surfactant proteins produced by the distal lung and secretoglobins and mucins from the airway (Fig. 3G). Notably, we discovered the secretion of both SCGB3A2 and SCGB1A1 peptides, which are specifically produced by a recently discovered human respiratory airway secretory (RAS) cell type^36^. This distinct cell type exists in the respiratory bronchioles region that is not found in mice and serves as unidirectional progenitors for type 2 alveolar cells to maintain the gas-exchange compartment. Functional proteins critical for lung development and homeostasis were also identified, including BPIFB1, A2M, and C6 involved in immune modulation, and collagen and fibronectin contributing to early lung structure formation and remodeling. In summary, the establishment of a perfusable human-specific capillary network in the subepithelial region of transplanted organoids offers a unique platform for investigating endothelial-epithelial crosstalk during organogenesis.

**Fig. 3.**
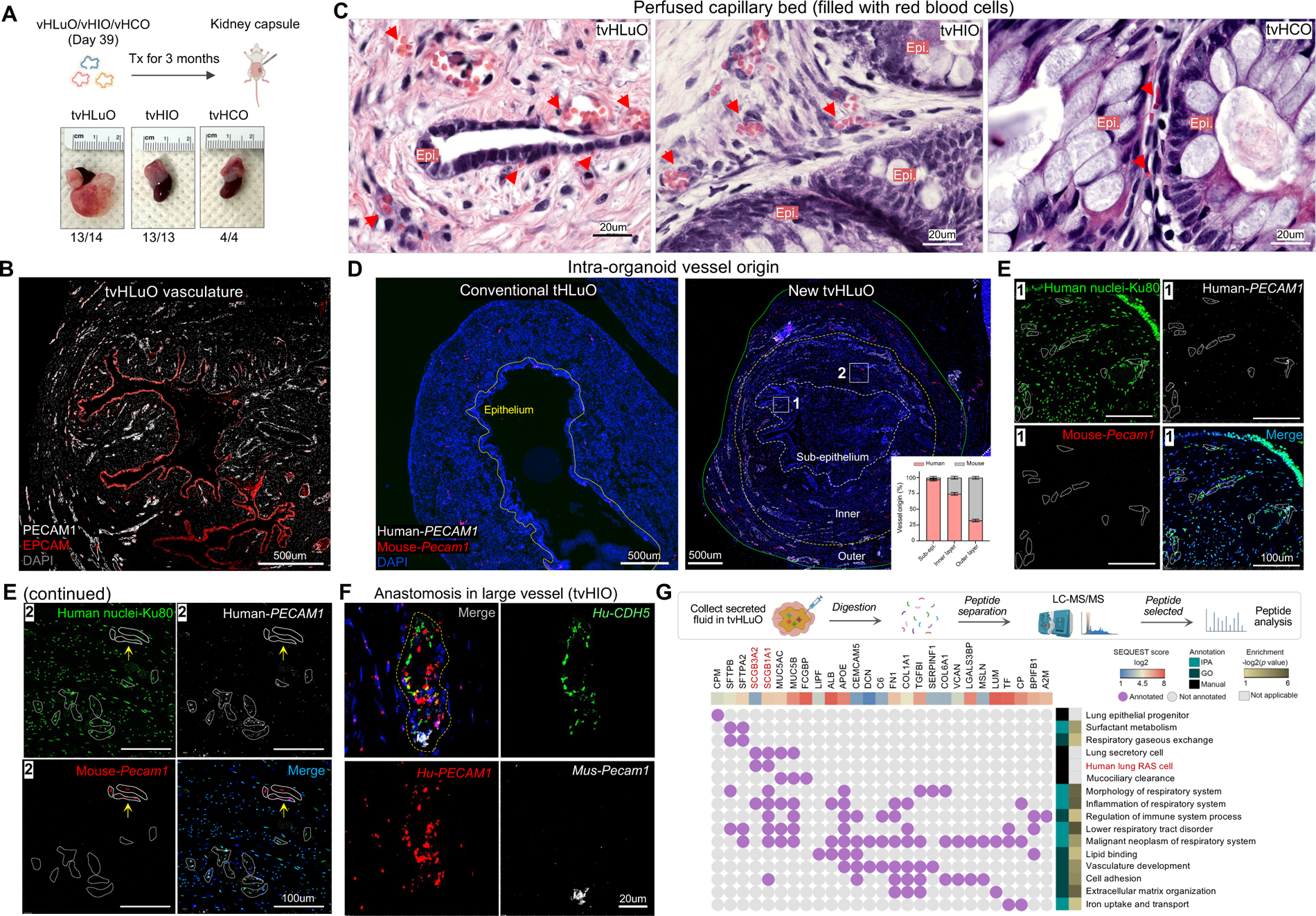
In vivo maturation of human vascularized lung and intestine organoid. **A.** In vitro vascularized organoids were further matured by transplantation under the kidney capsule of immune-compromised mice for three months. The numbers indicated the successful vs. total transplanted organoid. **B.** Immunostaining image of endothelial cells (PECAM1) and epithelial cells (EPCAM) within tvHLuOs. **C.** H&E staining of the explanted organoids showed sub-epithelial capillary beds filled with red blood cells (RBCs). Arrowhead: capillary vessel filled with RBC. **D.** Detecting the origin of vessels within conventional (left) and vascularized lung organoids (right) by human-specific and mouse-specific *PECAM1* smFISH probes and Ku80 human nuclei antibody. Dashed lines circled different layer structures based on the distance from the epithelial layer. Insert: Quantification of vessel origin within tvHLuO. Mean ± SEM. n= 10 regions from two different tvHLuO samples. **E**. Zoom in images of zone 1 (sub-epi) and zone 2 (inner layer) in panel D. **F.** Anastomosis structure connecting human and mouse vessels in tvHIOs. **G.** Proteomic analysis of fluid combined from two tvHLuO after 3-month transplantation. The secreted proteins within tvHLuO fluid were collected, digested with Trypsin, and analyzed by nano LC-MS/MS. The functions of identified protein peptides were further analyzed by either manual annotation based on known knowledge, or Gene Ontology (GO) and IPA pathway enrichment. The confidence of peptide identification is evaluated by SEQUEST score. The red color highlighted the human lung RAS cell markers. See also Fig. S5.

### Distinct organotypic features in vascularized gut organoid mesenchyme

To uncover regional patterns and distinctive organotypic features within the vascularized organoids, we conducted comprehensive computational analyses using scRNA-seq data from day-7 vAFG and vMHG, as well as day-21 vHLPO, vHIO, and vHCO organoids. Firstly, we re-analyzed the scRNA-seq dataset derived from the E8.75 mouse primitive gut tube, which included separately barcoded cells from anterior versus posterior primitive gut^34^ (Fig. 4A, Fig. S6A-S6B, and Table S2). Interestingly, we observed A-P patterning predominantly in the mesenchyme and epithelium, with less pronounced effects in the ECs (Fig. 4B). To determine the A-P patterning of the day-7 vAFG and vMHG organoids, we performed pseudo-bulk similarity analysis and demonstrated a positive correlation between human organoids and mouse embryonic tissues regarding the A-P patterning of the mesenchyme and epithelium. However, there was no discernible A-P difference observed in the ECs (Fig. 4C and Fig. S6C). Additionally, our scRNA-seq analysis unveiled both known (*Hoxb6*, *Hand1*)^9^ and novel markers distinguishing the anterior from posterior gut tube mesenchyme (Fig. 4D and Table S3). Several new genes shared by both human and mouse, such as *IRX3*, *IGFBP5*, *NR2F1*, *SFRP1,* and *MSX1*, were preferentially expressed in either anterior or posterior primitive gut tube organoid mesenchyme; multiple novel genes (*FOXC1*, *PRRX1*, and *HAPLN1*) were also discovered in human datasets that were exclusively expressed in the corresponding gut mesenchymal regions. Their expression was subsequently validated in human vAFG and vMHG organoids (Fig. 4E and Fig. S6D).

**Fig. 4.**
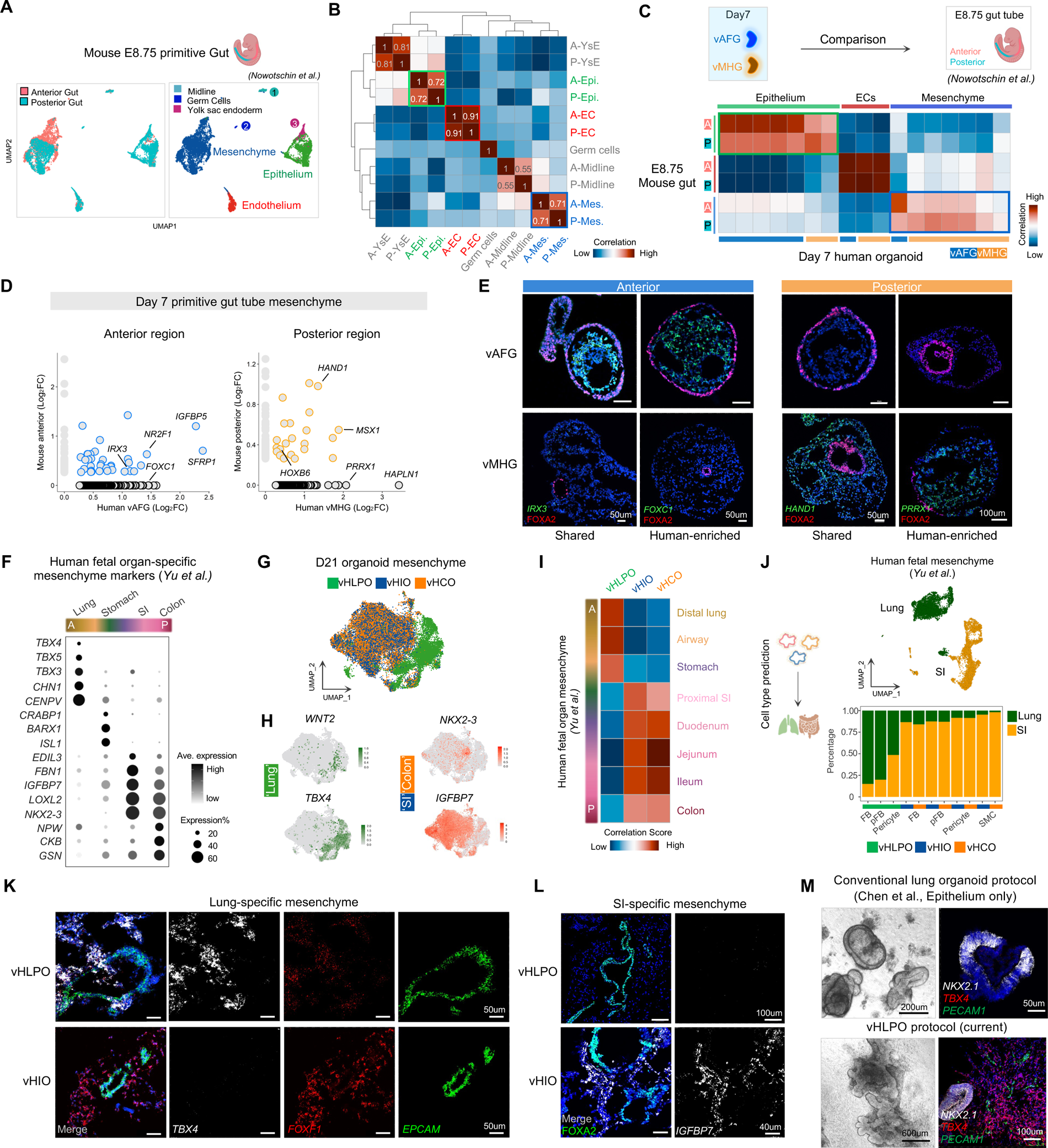
Characterization of the organotypic mesenchyme within vascularized organoids. **A.** UMAP projection of scRNA-seq datasets from E8.75 mouse anterior and posterior primitive gut tube (Nowotschin et al.,). **B.** Pseudo-bulk similarity analysis of major cell types within anterior versus posterior E8.75 mouse primitive gut tube. Mesenchyme and epithelium showed distinct A-P regional features. YsE, yolk sac endoderm; EC, endothelial cell. Mes: Mesenchyme; A, anterior; P, posterior. **C.** Pseudo-bulk similarity analysis between human vascularized organoids and E8.75 mouse gut. Both epithelium and mesenchyme exerted a high A-P pattern correlation between human organoids and mouse primitive gut. Colors represent the similarity correlation between human and mouse subclusters. **D.** scRNA-seq identified anterior- and posterior-selective genes of mouse and human primitive gut mesenchyme. **E.** smFISH revealed anterior and posterior mesenchymal genes in day 7 vascularized primitive gut organoids. **F.** Dot plot showing marker genes of human fetal organ-specific mesenchyme based on scRNA-seq data reported by Yu et al.. A: anterior; P: posterior. SI: small intestine. **G.** UMAP projection of mesenchymal cells from vHLPO, vHIO, and vHCO with Harmony integration. **H.** Feature plots of lung- and SI/colon-specific mesenchymal marker genes. **I.** Pseudo-bulk similarity analysis on mesenchyme between human fetal organs and vascularized organoids. SI: small intestine. **J.** Automated cell annotation of organoid mesenchymal populations based on human fetal lung and SI mesenchyme reference map (Yu et al.,). The predicted fetal lung and SI identities in each organoid mesenchymal cell type were normalized as percentile. **K-L.** smFISH analyses showing organotypic mesenchymal markers identified in **F.** in day-21 vHLPOs and vHIOs. *FOXF1,* generic mesenchyme; *EPCAM/FOXA2*, epithelium. **M.** Brightview and smFISH images of epithelium (*NKX2.1*), mesenchyme (*TBX4*) and endothelium (*PECAM1*) in day-21 conventional lung organoids and current vHLPOs. See also Fig. S6 and Table S2-S5.

To further determine the organotypic signatures of the mesenchyme within day-21 vascularized organoids, we analyzed scRNA-seq data of the mesenchymal cells from human fetal endodermal organs^35^ (Fig. 4F), and combined the mesenchymal populations from day-21 vHLPO, vHIO, and vHCO using Harmony-based integration (Fig. 4G). Intriguingly, the mesenchymal populations of lung and intestinal organoids were distinctly segregated and expressed corresponding human fetal gut mesenchymal markers: Lung mesenchymal cells expressed high levels of *WNT2* and *TBX4*^37^, whereas intestinal mesenchyme specifically expressed *NKX2-3* and *IGFBP7 (*Fig. 4H and Table S4-5). Additionally, pseudo-bulk analysis revealed the similarity of vascularized organoids to human fetal gut tissues regarding their mesenchymal properties along the A-P axis (Fig. 4I and Fig. 6E). This similarity was further confirmed via predicting the major organoid mesenchymal populations as those found in fetal organs (Fig. 4J). The organotypic features of the vHLPO and vHIO were further validated through smFISH staining of the lung-specific marker *TBX4* and the intestine-specific marker *IGFBP7*, while the early pan-splanchnic mesoderm marker *FOXF1* showed no difference (Fig. 4K-4L). Collectively, compared with conventional lung organoids^23^ (Fig. 4M, upper panels), the current vascularized organoid model (Fig. 4M, lower panels) showed complex epithelial morphology and contained organotypic mesenchyme (*TBX4*+ for lung mesenchyme^37^).

### Interactions among organotypic mesenchyme and neighboring cells

To uncover the key signaling contributing to the fate determination of organotypic mesenchyme within vascularized organoids and human fetal gut tissues, we performed CellChat analysis based on scRNA-seq. The ligand-receptor (L-R) pairs involved in communications from other cell types to mesenchyme were determined (Fig. S6F and Table S6-S7). Our findings revealed the expression of WNT7B within vHLPO epithelium, which targets FZDs and LRP6 expressed by lung mesenchyme^38^ (Fig. S6F). We also identified several L-R pairs shared by both vascularized organoids and human fetal tissues that could potentially drive the differentiation of organotypic mesenchyme *in vitro* (Fig. S6G, pink box). For example, GDF15 secreted by the intestinal epithelium may promote differentiation of posterior intestinal fibroblasts by binding to TGFBR2 receptor; FGF9 expression was observed in the lung epithelium, contributing to the specification of lung mesenchyme. Moreover, the vasculature within the organoids exerted an influence on mesenchymal patterning through the secretion of specific morphogens, including PROS1, SEMA6B, and EFNA3, which contributed to the development of intestinal mesenchyme.

### Organotypic endothelium in vascularized lung and intestine organoids

Compared to arterial or venous ECs, capillary ECs preserved the most organotypic features across different tissues^39^. To evaluate the organ specificity of capillary ECs in vascularized lung and intestine organoids, we reanalyzed multiple independent human cell atlases^4,5,35,40^. We identified specific markers for ECs in human lung, esophagus, stomach, small intestine (SI), and colon (Fig. 5A, Fig. S7A-S7B, and Table S8-S9). Intriguingly, upon selection and Harmony-based integration of EC clusters from vHLPO, vHIO, and vHCO, distinct cell distributions were observed (Fig. 5B): lung ECs clearly segregated from the intestinal ECs. Additionally, these ECs within the organoids expressed corresponding marker genes of human fetal lung (*HPGD, APCDD1*), SI (*STAB1, MTUS1*), and colon EC (*THSD7A, CXXC5*) (Fig. 5B, Fig. S7C, and Table S10-S11).

**Fig. 5.**
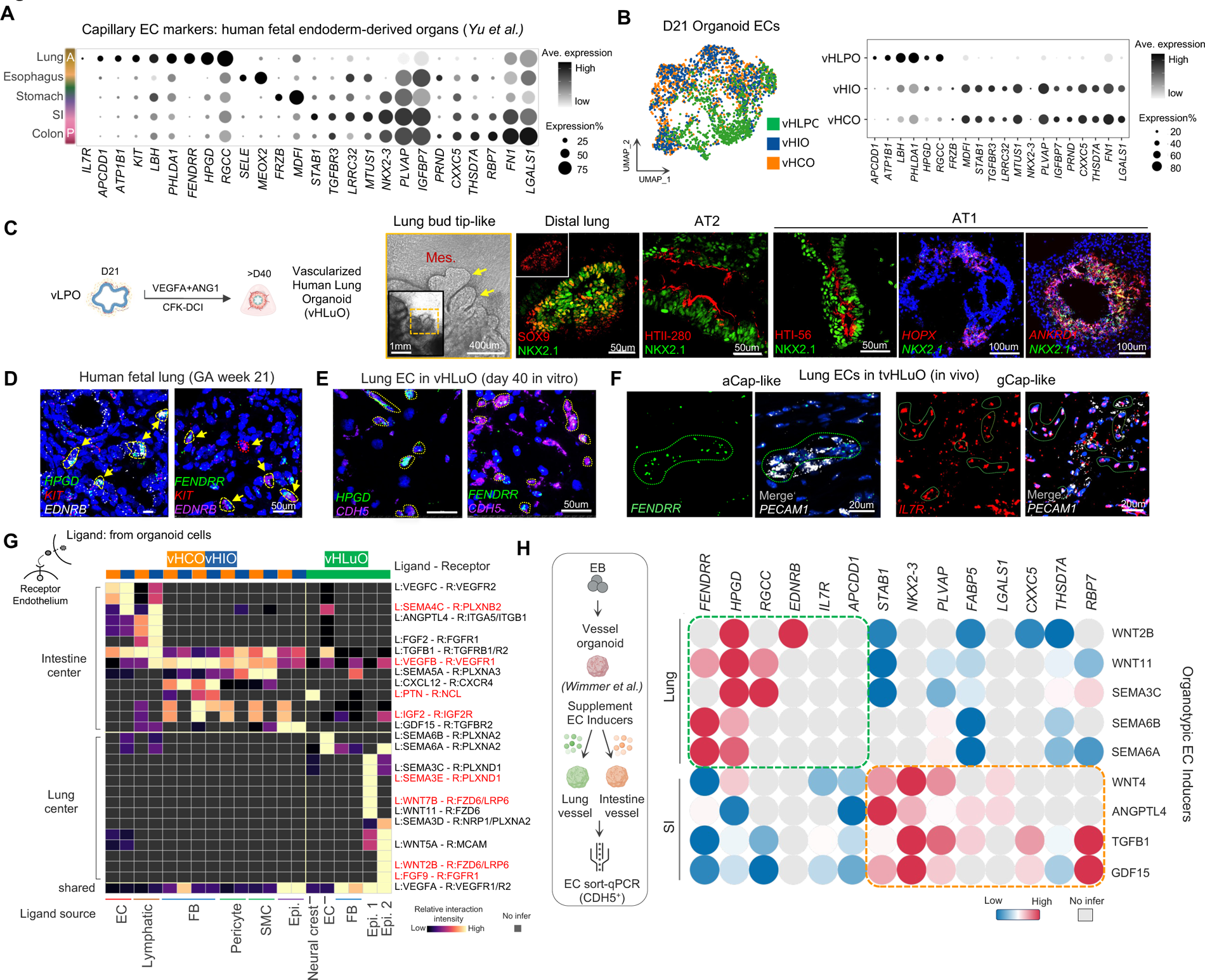
Determining morphogens for driving organotypic endothelial cell fate. **A.** Dot plot of organotypic EC markers generated based on scRNA-seq datesets of human fetal organs published by by Yu et al. EC, endothelium. SI, small intestine. **B.** UMAP projection and dot plots of organ-specific EC markers in day 21 vHLPOs, vHIOs, and vHCOs. **C.** Brightview image (left) showing lung bud tip-like morphology (yellow arrows) observed in day-40 vHLuOs. Immunostaing of day-40 vHLuOs (right) showing distal lung epithelial marker (SOX9, NKX2.1), AT2 marker (HTII-280), and AT1 marker (HT1-56). Additional AT1 markers were determined by smFISH (*HOPX*, *ANKRD1*). **D.** smFISH detection of lung EC markers in human fetal lung tissues. Yellow arrows: lung-specific ECs. GA: gestational age. **E-F.** Detection of human lung EC markers in day-40 vHLuO or transplanted tvHLuO. *PECAM1* and *CDH5* were used as pan-EC markers. **G.** CellChat analysis of vascularized organoid scRNA-seq unveiled key ligand-receptor pairs orchestrating cell-cell communication. The heatmap highlighted receptors expressed by ECs, with corresponding ligands secreted by neighboring cell types. The dark gray blocks indicated L-R pairs undetected. L, ligand; R, receptor. The red-highlighted ligand-receptor pairs denote those shared between vascularized organoids and human fetal tissues. **H.** To induce organotypic EC specification, key ligands revealed in panel G were supplemented to the differentiation medium of generic vessel organoids from day 5 to day 15. Samples were collected on day 15 and qPCR analysis of organ-specific EC marker genes were performed in ECs isolated from the vessel organoids. The qPCR results were summarized in the heatmap. N=3 repeats in each treatment group. See also Fig. S7 and Table S6-S11.

Next, to generate highly specialized alveolar capillary ECs mediating gas exchange^5,40^, we further differentiated day-21 vHLPO into a more mature vascularized human lung organoid (vHLuO). By day 40, vHLuO expressed marker genes of distal lung epithelium (SOX9^+^/NKX2.1^+^), AT2 cells (HTII-280), and AT1 cells (HT1-56/*HOPX*/*ANKRD1*)^26^ (Fig. 5C). To characterize the alveolar capillary ECs, firstly we validated the scRNA-seq data and demonstrated the enriched expression of lung EC markers (*HPGD* and *FENDRR*)^40^ in human fetal lung tissue at Gestational Age (GA) week 21 (Fig. 5D) and postnatal human lung (Fig. S7D), while absent in other examined organs (Fig. S7E). Consistently, lung EC-specific gene *HPGD* was also expressed in day 21 vHLPO (Fig. S7F) and day 40 vHLuO (Fig. 5E), but not vHIO (Fig. S7G). Conversely, the expression of SI/colon EC marker *IGFBP7* was enriched in vHIO compared to vHLPO (Fig. S7H).

To further characterize the recently discovered alveolar capillary EC subtypes, known as aerocyte (aCap) and general capillary ECs (gCap) ^5,40^, we analyzed vHLuOs three months post-transplantation (tvHLuO). The smFISH analyses demonstrated the presence of both aCap-like cells (*FENDRR*^+^) (Fig. 5F) and gCap-like cells (gCap, *IL-7R*^+^/*KIT*^+^) (Fig. 5F and Fig. S7I). Collectively, the vascularized organoids contained not only organotypic mesenchyme and epithelium, but also different subtypes of the endothelium such as aCap- and gCap-like alveolar ECs.

### Interactions among organotypic endothelium and neighboring cells

To further determine the specific ligands that drive the development of organ-specific ECs, CellChat analysis was performed based on scRNA-seq of day-21 vascularized organoids (Fig. 5G and Table S6) and human fetal tissues (Fig. S7J and Table S7). In the lung, WNT2B expressed by mesothelium and WNT7B expressed by epithelium interacted with FZD6/LRP6 expressed by the lung ECs, which is consistent with previous findings^38,41^. In the intestine, VEGFB expressed by various mesenchymal cells such as fibroblasts and pericytes, bound to the VEGFR1 expressed by the endothelium (Fig. 5G)^42^. Several ligands targeting ECs were commonly identified in both organoid and human fetal tissues and have not been demonstrated as organotypic EC drivers previously, such as VEGFB (from intestine epithelium), IGF2 (from intestine epithelium and mesenchyme), FGF9 (from lung mesenchyme), and SEMA3E (from lung epithelium) (Fig. 5G and Fig. S7J). Next, we assessed the capacity of these ligands in patterning organ-specific ECs in generic 3D vessel organoids (VOs)^43^. The ECs within the generic VOs lack distinct organ-specific characteristics, as demonstrated by scRNA-seq analysis, cross-referencing with the human fetal organ atlas, and pathway enrichment analysis^44^. After differentiating the VOs through early mesoderm and angioblast progenitor stages, we exposed the day 5 VOs to candidate morphogens individually for another 10 days to generate organotypic ECs. On day 15, CD31^+^ ECs from VOs were enriched for the assessment of organ-specific EC features at the transcriptomic level. We found that WNT2B, WNT11, SEMA3C, SEMA6A, and SEMA6B were effective in upregulating the expression of lung-specific marker genes in VOs, whereas WNT4, ANGPTL4, TGFB1, GDF15 were effective in driving generic ECs towards an intestinal cell fate (Fig. 5H and Fig. S7K). These findings underscore the pivotal role of the microenvironment in directing and maintaining the organ-specific characteristics of ECs.

### Modeling congenital lung defects caused by FOXF1 variants in vHLuO

Compared to conventional lung epithelial organoids, our current multilineage lung organoid model holds an advantage due to its incorporation of FOXF1-positive mesenchymal and endothelial populations. Consequently, we aimed to leverage this model for investigating a severe lung developmental disorder stemming from variants involving the *FOXF1* gene locus - Alveolar Capillary Dysplasia with Misalignment of Pulmonary Veins (ACDMPV, or ACD). Although the *Foxf1* mutant mouse model has been created to study ACD^45,46^, there are variations in gene signatures of EC subtypes and developmental timing of the alveolarization in mouse vs. human, limiting its application in fully recapitulating human pathology and understanding ACD etiology.

We generated vHLuOs from iPSCs derived from three ACD patients and three healthy controls (Fig. 6A and Fig. S8A). One patient (ACD-1, also known as ‘FOXF1.1’ iPSC line) has a 1.7Mb chromosome deletion containing the *FOXF1* locus^47^, while the other two carried different heterozygous *FOXF1* mutations (c.166C>G and c.253T>A, ACD-2 and ACD-3)^48^ (Table S12). We observed significantly decreased capillary formation (Fig. 6B) and reduced aCap and gCap populations in all three ACD vHLuOs compared with controls (Fig. 6C-D). Interestingly, despite the absence of FOXF1 expression in lung epithelial cells, we noted a decrease of distal lung epithelium (SOX9^+^, NKX2-1^+^) (Fig. 6E), impaired AT1 differentiation (*HOPX*^+^) (Fig. 6F), and an accumulation of damage-associated AT1/AT2 intermediate cells (*CLDN4*^+^/*NKX2.1*^+^) (Fig. 6G) in ACD vHLuOs^49–51^. These cell non-autonomous abnormalities in epithelial cells are likely attributed to the surrounding defective endothelial and mesenchymal populations and disrupted cell-cell crosstalk. Notably, these cellular phenotypes closely resembled those found in human ACD lung tissues^52^ (Fig. 6K), and could not be fully replicated in the *Foxf1*-mutant mouse model^45^ or traditional lung organoid model due to its absence of mesenchymal and endothelial components (Fig. 6H-6J and Fig. S8B-S8C).

**Fig. 6.**
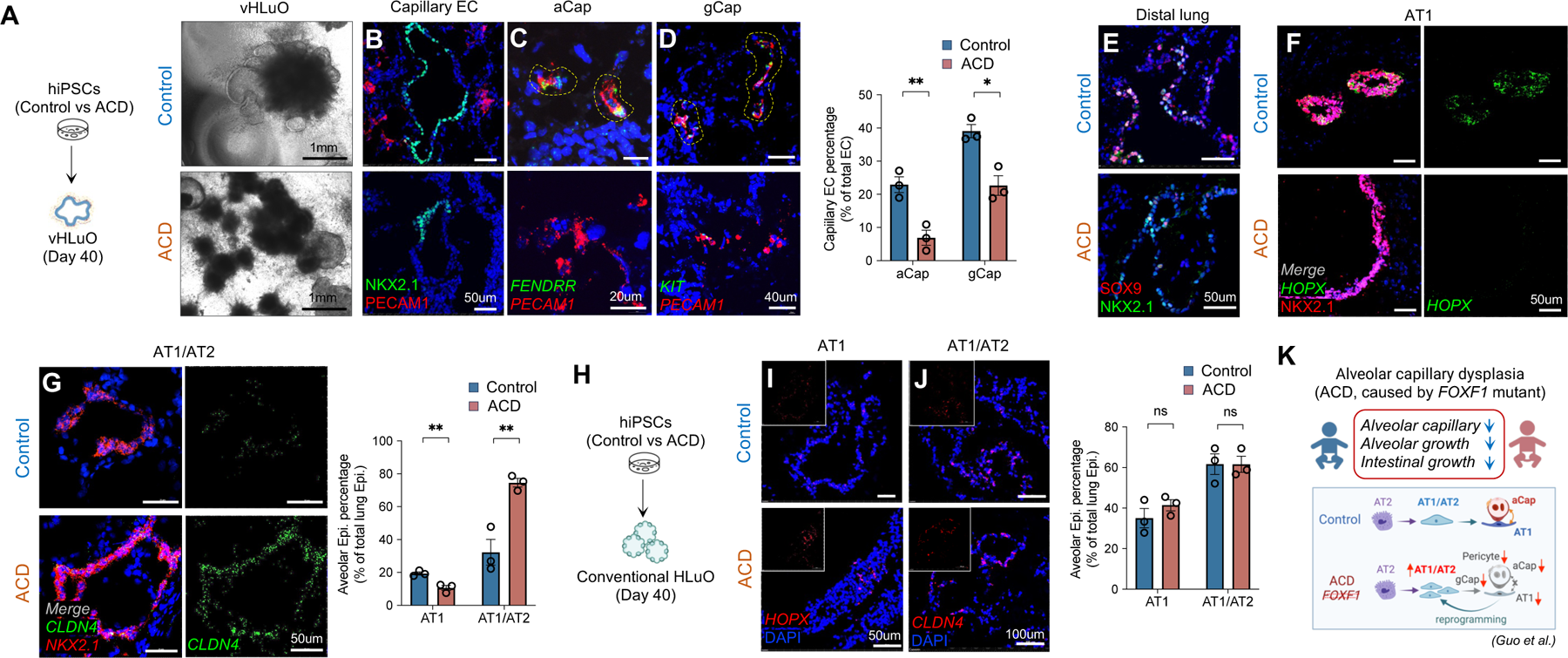
Determining cellular phenotypes caused by *FOXF1* mutations utilizing patient-specific vHLuOs. **A.** Bright-field images of day-40 vHLuO differentiated from healthy control or ACD iPSCs. **C-D.** Immunostaining of lung capillary EC markers in both control and ACD vHLuOs and statistical quantification of the percentage of capillary EC subtypes. Yellow dashed lines highlighted the *FENDRR^+^/PECAM1^+^* aCap or *KIT^+^/PECAM1^+^* gCap. **E-G.** Immunostaining for distal lung, AT1, and AT1/AT2 markers in control and ACD vHLuOs, followed by statistical quantification of the percentage of alveolar epithelial subtypes. **H-J.** Immunostaining of AT1 and AT1/AT2 markers in both control and ACD conventional HLuOs followed by statistical quantification. **K.** Cellular abnormalities attributed to FOXF1 mutations identified through scRNA-seq analysis of lung tissues from control and ACD patients, as reported by Guo et al. All data represent mean ± SEM. *p<0.05, **p<0.01, ACD vs. Control. Unpaired t-test. N= 3 different biological samples. See also Fig. S8 and Table S12.

### Modeling intestinal defects caused by FOXF1 variants in vHIOs

In addition to the lung abnormalities, ACD patients often have extrapulmonary anomalies, such as small intestinal atresia and malrotation^53^. Our current vHLuO and vHIO organoid models provide a unique opportunity to comprehensively study both pulmonary and intestinal defects in ACD patients. In three ACD vHIOs, we observed reduced endothelial (PECAM1^+^) and mesenchymal (FOXF1^+^) populations compared to controls (Fig. S8E-S8F). The mesothelial population (WT1^+^) was also found to be suppressed in ACD vHIOs (Fig. S8F), which is crucial for the formation of connective tissues that facilitate intestinal movement and suspension^54^, and a major source of smooth muscle cells in the gut vasculature^55^. The reduction of mesothelial cells might offer a partial explanation for the intestinal malrotation observed in ACD patients. Additionally, ACD vHIOs also showed an impaired differentiation of posterior gut tube epithelium (FOXA2^+^, CDX2^+^) alongside the appearance of ectopic anterior marker (SOX2^+^) (Fig. S8D and S8G).

In summary, our study demonstrated that the current organoid system is superior in deciphering intricate cell-cell communication and uncovering both cell-autonomous and non-autonomous phenotypes. This offers a unique platform to investigate complex genetic disorders involving multiple cell types in both pulmonary and intestinal organ systems.

## Discussion

In our study, we effectively engineered vascularized lung and intestinal organoids from hPSCs with organotypic endothelium and mesenchyme by precisely controlling of mesoderm-endoderm co-differentiation and inducing specific primitive gut tube lineages. Upon transplantation under the kidney capsule in mice, these multilineage organoids, rich in human-specific vasculature, integrated with the host circulation, fostering further growth and maturation. This strategy addressed the limitations of existing models^56,57^ by generating a diverse array of cell types originating from both mesoderm and endoderm, such as aerocytes^40^ and RAS cells^36^, which have not been described in previous lung organoid models yet crucial for human alveoli development and regeneration. The cell type composition within the multilineage organoids closely resembled that of cells found in human fetal organs. It also recapitulated the concurrent development of multiple germ layers, fostering essential cell-cell communication during the initial phase of human development-an aspect lacking in assembloid models. Additionally, it provided a unique system for defining marker genes that distinguish anterior versus posterior gut tube mesenchyme, which had previously remained unclear due to the restricted availability of early human embryonic tissues. More importantly, our study identified novel ligand-receptor pairs responsible for the cell fate determination of organ-specific endothelium and mesenchyme.

Our research uncovered the significant impact of BMP signaling during the initial three days of differentiation in regulating the balance between mesoderm and endoderm, and in pre-determining the anterior-posterior patterning of the gut tube epithelium. While previous studies have primarily focused on inducing either DE or mesoderm lineages via the regulation of TGFb and Wnt/BMP pathways ^6,27,28,43^, our approach aimed to recapitulate human embryonic development and create spheroids comprising both DE and mesoderm cells. We observed that while Wnt activation was indispensable to prime the meso-endodermal differentiation^12,13^, controlling the duration of BMP activation was critical to fine-tune the ratio of DE and mesoderm within a single spheroid. More interestingly, we found that manipulating BMP signal activity during the first three days of differentiation resulted in heterogenous DE subclusters in the meso-endodermal spheroids, predetermined the anterior-posterior patterning of the gut tube epithelium in the subsequent organogenesis phase (day 7-21). The diverse DE clusters observed aligns with findings reported by Mahaddalkar et al., which identified multiple gut tube progenitors potentially contributing to the production of more mature insulin-producing b-like cells^58^. While this finding broadly corresponds with the recognized role of BMP signaling in embryonic posteriorization^7^, it unveils an unexpected predisposed gut tube lineage fate pre-determined by BMP4 signaling as early as at the DE stage.

Organ-specific mesenchyme significantly contributes to organ development and regeneration, yet previous endeavors predominantly focused on differentiating gut tube epithelial organoids, often with a poorly characterized and limited amount of accompanying mesenchyme^27,28,59,60^. In a 2D context, Han et al.^9^ and Kishimoto et al.^61^ established an iPSC-based roadmap for generating foregut organ-specific mesoderm based on signaling networks and molecular markers elucidated from single-cell transcriptomics of mouse foregut organogenesis. Building upon this, Eicher et al. utilized these iPSC-derived splanchnic mesoderm cells to generate human gastric organoid containing the smooth muscle layer^11^. Alber et al. identified the crucial role of retinoic acid and hedgehog signals in specifying early lung mesenchyme positive for TBX4, facilitating the generation of functional mouse iPSC-derived lung mesenchyme using reporter lines^62^. However, existing strategies rely on cell sorting using engineered reporter lines, or lack the intricate 3D architecture necessary for effective tissue modeling. In contrast, our system successfully generated lung- and intestinal-specific mesenchyme within organoids that were benchmarked against corresponding human fetal mesenchymal cells through single-cell RNA-seq. Additionally, these cells self-organized alongside epithelial cells, creating a microenvironment for investigating cell-cell interactions during organ formation. More importantly, our study unveiled a range of growth factors secreted by the neighboring cells within the organoids critical in guiding the specification of the tissue-specific mesenchyme. For instance, beyond the previously reported BMP2^27^, we identified a potential role of GDF15 in promoting intestinal mesenchyme specification. These signaling pathways hold promise for direct differentiation of tissue-specific mesenchyme from iPSCs in future applications.

To vascularize organoids, we also assessed the optimal developmental timing for introducing vasculogenesis factors. Prior to gut tube formation, vessels begin developing in the splanchnic mesoderm within the embryo, ultilmately contributing to the primary vasculature and capillary plexus in the gut wall^63^. During the formation of primitive gut tube in mouse embryos (E8.25), ECs are autonomously generated in the yolk sac, allantois, and notably, the embryo proper^18^. In our human organoid system, we also observed spontaneous emergence of the vascular bed within the splanchnic mesoderm in day-7 vAFG and vMHG, which was further expanded and matured until day 21 through supplementation with Angiopoietin 1^25,63^. Interestingly, contrary to the vMHG response, the introduction of additional VEGFA during vAFG differentiation unexpectedly inhibited the specification of anterior lung epithelial progenitors. Previous studies have suggested that VEGF activation could induce BMP signaling during mesendodermal patterning^64^, potentially imprinting posterior features into the vAFG and resulting in the observed mispatterning event. These findings underscore the importance of precise temporal control of angiogenic signals when recapitulating vasculogenesis in different organs, and emphasize the intricate and varied nature of the vascular emergence process across different organ systems.

Vascularization of the organoids, particularly for the lung, continues to pose a significant technical bottleneck. Previous studies by Holloway et al.^29^ and Childs et al.^65^ introduced angiogenic factors—VEGFA, BMP4, and FGF2—or Epiregulin (EREG) during the generation of small intestinal organoids to enhance the growth of endogenous vessels. While this approach generated ECs that partially resembled the human fetal small intestine endothelial gene profile, the robustness of the vasculature fell short of that observed in the human fetal counterpart and in *in vivo* transplantation models, which was predominantly comprised of invaded mouse vessels. More importantly, current methods for generating lung organoids have yet to demonstrate the formation of the human pulmonary vasculature, potentially attributed to prior protocols emphasizing anterior foregut generation, necessitating strong BMP and TGFb inhibition which could have hampered vasculogenesis. Through implementing BMP inhibition at optimal levels, meticulous adjusting the ratio between mesoderm and endoderm, and precise temporal control of angiogenic cues, we successfully introduced human vasculature with organ specificity into lung, small intestine, and colon organoids. Remarkably, these vessels retained their human and organ specificity for three months post-transplantation under the mouse kidney capsule. These vessels also integrated with the host circulation as demonstrated by the perfusion of red blood cells within the human vasculature. Intriguingly, nearly all capillary networks growing adjacent to the epithelial layers originated from the transplanted human organoid with minimal regression or penetration issues, which stands in contrast to the challenges consistently observed in previous studies that host-derived vasculature hindered the advanced maturation of transplanted organoids^23,29^.

Another significant challenge in the field is the generation of organotypic ECs that faithfully replicate primary tissue-specific ECs at both molecular and functional levels, particularly those relevant to gut organs. Herein, through analyzing scRNA-seq datasets from our vascularized organoids and publicly available human fetal gut organs, we consolidated a list of novel morphogens that specifically target lung or intestinal ECs. Subsequently, we evaluated the effectiveness of these newly identified morphogens in driving the differentiation of organotypic ECs from iPSCs-derived vessel organoids that contain generic EC progenitors. Wnt family gene expressions are found in both gut epithelium and mesenchyme and correlate with mouse embryonic gut regional specification. Among the various Wnt family genes, Wnt4 was found in the posterior intestinal epithelium, while Wnt11 is expressed in the anterior esophagus and stomach^66^. Apart from controlling gut epithelium patterning, we found human WNT4 ligand also enhanced the expression of genes such as *STAB1*, *NKX2-3*, and *PLVAP*, which were representative of intestinal EC features. On the other hand, WNT11 induced the expression of lung EC (*RGCC*) and aerocyte genes (*FENDRR* and *HPGD*) while suppressing the posterior intestinal EC gene signature. Semaphorin signaling is involved in multiple processes contributing to circulatory system development and emerging data highlights the crucial role of Semaphorin in the development of the kidney and lung regarding their branching morphogenesis^67^. Likewise, we showed that certain Semaphorins were effective in inducing lung EC over intestinal EC gene signatures. Collectively, these novel EC morphogens hold great potential to yield specialized organotypic vasculature with high efficiency.

In addition to study human organ development, the current multilineage vascularized organoid model provided a suitable venue for recapitulating complex genetic disorders, particularly in uncovering cell non-autonomous mechanisms. Here, we showcased that severe underdevelopment of pulmonary and intestinal epithelium in ACDMPV is directly attributed to aberrant vascular and mesenchymal populations driven by the *FOXF1* loss-of-function mutation. However, the conventional lung organoid models, limited by their absence of FOXF1-expressing cells mediating multicellular interactions, failed to recapitulate the epithelial abnormalities and establish comprehensive genotype-phenotype connections. Our multilineage organoids provide a unique platform to investigate both cell autonomous and non-autonomous phenotypes in a variety of congenital lung disorders, such as pulmonary hypoplasia and alveolar dysplasia caused by *TBX4* mutation^68^, a key transcription factor exclusively expressed in the lung mesenchymal compartment.

In this study, we present an efficient platform to generate organotypic mesenchyme and vasculature by orchestrating mesoderm-endoderm co-differentiation guided by embryogenesis and vasculogenic cues. Notably, this approach is not confined to lung or intestine organoids, but can be adapted to create a variety of multilineage organoids featuring both vascular and mesenchymal compartments, broadening its applicability to other organs such as the stomach and pancreas. It is important to note that the cells within these organoids exhibit characteristics akin to the human fetal stage, suggesting that achieving advanced functional capabilities may necessitate further maturation of the system. Given the intricate 3D architecture and cellular complexity of the iPSC-derived vascularized organoids, they present a promising avenue for personalized organoid therapy, especially in organs with limited self-regeneration capacity. Additionally, as ECs are the primary targets of the immune system during organ transplantation rejection, vascularized organoids offer immense potential in unraveling the mechanisms underlying EC-mediated allograft failure in antibody-mediated rejection.

## Acknowledgments

We thank the Discover Together Biobank for support of this study, as well as the participants and their families, whose help and participation made this work possible; Dr. Matt Kofron from Confocal Imaging Core (CIC), CCHMC, and Joseph Kitzmiller from Division of Pulmonary Biology, CCHMC for providing access and assistance to the confocal microscope and image processing. Mass spectrometry data were collected and analyzed in the University of Cincinnati Proteomics Laboratory under the direction of Ken Greis, Ph.D. on instrumentation supported by an NIH high-end instrumentation grant (S10OD026717-01). This work was supported by CCRF Endowed Scholar Award and R01HL166283 (M. Gu). N.M.P received the American Heart Association Pre-Doctoral Fellowship grant 1013861. Z.Y received the American Heart Association Pre-Doctoral Fellowship grant 906513. T. T received the Falk Transformational Awards Program and NIH DP2 grant DK128799-01. R.J.R received Human Disease Model Award of the Erasmus MC. D.N.K received NHLBI/NIH grants: N01-75N92020C00005 and R01HL095993.

## Author Contributions

Y.M. and C.T. contributed equally to this work. Y. M., C.T., M. Guo, and M. Gu conceived and designed experiments. N.M.P. performed the VO in vitro assay. Y. M., Z. Y., D.O.K., M.C.Y., and Y.W.C. optimized and generated organoids. C.J. performed the bioinformatic analysis supervised by M. Guo. C.T., N.S., and Y.W.C. performed in vivo organoid transplantation. K. I., D.O.K., V. P. G., and K. K. performed staining and imaging analysis. J. T., J.A.W., K.C.M., R.J.R., D.N.K., M.A.H., J.M.W., T. T., and A.M.Z. contributed to all necessary iPSC cell lines, animals, material, and intellectual discussions. Y.M., C.T., N.M.P., Z.Y., D.O.K., N.S., M.C.Y., Y.W.C., M. Guo, and M. Gu wrote the manuscript with the contributions from all other authors. Y.M., M. Guo, and M. Gu oversaw the project. All authors read and approved the manuscript.

## STAR Methods

### RESOURCE AVAILABILITY

#### Lead contact

Further information and requests for resources and reagents should be directed to and will be fulfilled by the lead contact, Mingxia Gu.

#### Materials availability

This study did not generate new unique reagents.

#### Data and code availability

- The raw and processed data from single-cell RNA sequencing in this study have been deposited with the Gene Expression Omnibus (https://www.ncbi.nlm.nih.gov/geo/). Accession numbers are listed in the key resources table. This paper analyzes existing, publicly available data. Related information for those datasets is listed in the key resources table. The mass spectrometry proteomics data have been deposited to the ProteomeXchange Consortium via the PRIDE partner repository (https://www.ebi.ac.uk/pride/). Accession numbers are listed in the key resources table.
- All original code has been deposited at https://github.com/MZGuo-lab/Vascularized_human_organoids and is publicly available as of the date of publication.
- Any additional information required to reanalyze the data reported in this paper is available from the lead contact upon request.

### EXPERIMENTAL MODEL AND STUDY PARTICIPANT DETAILS

#### Human ESC/iPSC generation and maintenance

All experiments involving the generation and differentiation of human PSC lines were performed with the approval of the Institutional Review Board (IRB) of Cincinnati Children’s Hospital Medical Center (CCHMC), Boston University, and the Daily Board of the Medical Ethics Committee (METC) Erasmus University Medical Center Rotterdam, The Netherlands. Control-1 iPSC line (BU3 NGST iPSC line, male, 32-year-old), carrying NKX2.1-GFP and SFTPC-tdTomato reporters, was obtained from prior studies as described^22^. The control-2 iPSC line, purchased from the California Institute of Regenerative Medicine (CIRM) iPSC Repository, is derived from fibroblasts from a female donor of one-year-old. Control-3 iPSC line was derived in-house by using 20-month female donor’s fibroblasts obtained from Biorepository for Investigation of Neonatal Diseases of the Lung (BRINDL). ACD-1 iPSC line, also known as ‘FOXF1.1’ iPSC line, was reprogrammed from fibroblasts from a newborn female donor at the Center for Regenerative Medicine, Boston University^47^. ACD-2 (‘EMC127i-A:ACD871C4’) and ACD-3 (‘EMC128i-A:ACD874C9’) iPSC lines were derived from fibroblasts from two newborn male donors at the Erasmus University Medical Center^48^. Fibroblast samples from healthy control and ACD patients were procured with documented informed consent. H9-GFP hESC line (female) was generated as described previously^69^. Human iPSCs and ESCs fed with StemMACS iPSC-brew XF culture medium were routinely cultured on the Cultrex-coated 6-well plate at 37°C and passaged with 0.5mM EDTA solution in PBS every 3-5 days. All human stem cells were routinely authenticated by testing the karyotype abnormality, short tandem repeat analysis, pluripotency marker expression, and mycoplasma contamination.

#### Animals

Immune-deficient NOD-SCID IL-2Rγnull (NSG) mice at the age of 8–16 weeks old were obtained from the Comprehensive Mouse and Cancer Core Facility at CCHMC or the Icahn School of Medicine at Mount Sinai and used in vascularized organoids and conventional HLuO, HIO, and HCO transplantation experiments. Both male and female mice were used in the study and we did not observe any gender effect on the results. Littermates were randomly assigned to experimental groups. All experiments were performed with the approval of the Institutional Animal Care and Use Committee (IACUC) of CCHMC and the Icahn School of Medicine at Mount Sinai.

#### Human fetal tissue

De-identified human fetal tissues from gestational age 21-38 weeks were collected and requested through Discover Together Biobank at Cincinnati Children’s Research Foundation (CCRF) under the approved IRB. No significant gender differences were observed in the current study.

### METHOD DETAILS

#### Generation of conventional and vascularized organoids

Human pluripotent stem cells were first dissociated into single cells with 0.5mM EDTA and filtered by 40nm Flowmi. Following the manufacturer’s instruction, 3D Embryonic bodies (EBs) containing 500∼1000 cells per EB were made with AggreWell 400 in the Aggregation media including 50uM Y-27632 (50ml aggregation media included 40ml knockOut DMEM/F12, 10ml knockOut serum replacement, 0.5ml GlutaMAX, 0.5ml NEAA, 35µl 1:100 b-Mercaptoethanol in PBS, 0.5ml Antibiotic-Antimycotic). After 24 hours, EBs were formed and washed with N2B27 basal media (500ml media included 250ml DMEM/F12, 250ml Neurobasal medium, 10ml B27 supplement, 5ml N2 supplement, 2.5ml GlutaMAX, 350ul 1:100 b-Mercaptoethanol in PBS, 5ml Antibiotic-Antimycotic). 1200 EBs can be evenly distributed into 3 wells of ultralow attachment 6-well plate containing 2-3ml medium and be gently shaken on the orbital shaker at 90rpm speed.

##### Generation of vascularized lung organoid

Several lung organoid and vessel organoid differentiation protocols were scrutinized to establish the vascularized lung organoid platform^22–24,29,43,59^. EBs were differentiated into meso-endodermal spheroid using 72 hours (Day 0-3) of Activin A (100ng/ml) in N2B27 basal medium and increasing concentration of defined FBS (Day 0-1, 0%; Day 1-2, 0.2%; Day 2-3, 2%). During the first 24 hours, CHIR99021 and BMP4 (30ng/ml) were included. CHIR at 3uM, 6uM, 9uM, and 12 uM concentrations were tested to optimize the differentiation. For most hiPSC and hESC lines, 12 uM of CHIR induced full meso-endodermal differentiation in EBs at Day 3. In some cases, FGF2 (20ng/ml) or PI3K inhibitor (PIK90, 1uM) were tested for the first 24 hours^6^. vAFG was subsequently generated by adding Noggin (100ng/ml) into N2B27 medium with 2% defined FBS for 96 hours (Day 3-7). Media was changed daily from day 0 to day 7. CHIR (2uM), FGF4 (500ng/ml), SB-431542 (10uM), or SAG (1uM) were tested for vAGF induction efficiency from day 3-7. On day 7, vAFG was collected and resuspended in 150ul Collagen-Matrigel mixture and transferred to 12-well transwell insert (0.4um) to solidify at 37°C incubator for 30 mins. The collagen solution was made following this order: 0.1N NaOH, 750ul; 100x DMEM, 313ul; 1M HEPEs, 63ul; 7.5% NaHCO_3_, 49ul; 10X GlutaMAX, 31ul; Ham’s F-12, 460ul; 3mg/ml PureCol Collagen I, 3.33ml. The Collagen-Matrigel mixture was made by mixing three portions of collagen solution with one portion of Matrigel. vAFG was incubated with lung specification media 1 from day 7-10 (100nM ATRA, 10ng/ml BMP4, 3uM CHIR, 10ng/ml FGF7; 10ng/ml FGF10; 100ng/ml VEGFA; 100ng/ml ANG1 in N2B27 basal media) and then switched to lung specification media 2 from day 10-21 (100nM ATRA, 3uM CHIR, 10ng/ml FGF7, 10ng/ml FGF10, 100ng/ml VEGFA, 100ng/ml ANG1 in N2B27 media) to generated vHLPO. 1ml culture media was added to the bottom chamber of the transwell to create an air-liquid interface (ALI) and changed every other day. To generate distal vHLuO containing AT2 cells, vHLPO was exposed to the distal lung specification medium from day 21-28 (3uM CHIR, 10ng/ml FGF7, 10ng/ml FGF10, 50nM Dexamethasone, 0.1mM cAMP, 0.1mM IBMX, 100ng/ml VEGFA, 100ng/ml ANG1 in N2B27 media). Distal vHLuO can be further patterned into AT1 cell-enriched lung organoids using AT1-specification media for another 10 days (10ng/ml FGF10, 10uM LATS-IN-1, 50nM Dexamethasone, 0.1mM cAMP, 0.1mM IBMX, 100ng/ml VEGFA, 100ng/ml ANG1 in N2B27 media)^26^.

##### Generation of vascularized small intestine and colon organoid

Similar to generating vascularized lung organoids, EBs were exposed to 100ng/ml Activin A (Day 0-3) and 12uM CHIR (Day 0-1), while BMP4 (30ng/ml) was added for 72 hours (Day 0-3) in N2B27 media with increasing concentration of defined FBS (0%, 0.2%, 2%). vMHG organoids were generated after 96 hours of CHIR (3uM), FGF4 (500ng/ml), and VEGFA (100ng/ml). Media was changed daily. On day 7, vMHG was embedded in the same matrix as vHLPO on the transwell insert to create ALI. Spheroids were patterned in EGF (100 ng/mL), Noggin (100ng/ml), R-spondin-1 (5%), VEGFA (100ng/ml), and ANG1 (100ng/ml) for vHIOs or EGF (100ng/ml), BMP2 (100 ng/mL), VEGFA (100ng/ml), and ANG1 (100ng/ml) for vHCOs for 72 hours (Day 7-10). Both vHIOs and vHCOs were maintained in EGF (100ng/ml), VEGFA (100ng/ml), and ANG1 (100ng/ml) from day 10 to day 35. Media was changed every other day.

##### Generation of conventional human lung, intestine, and colon organoids

The generation of conventional lung organoids was performed based on previous studies^23,59^. Briefly, hPSC differentiation into endoderm was performed in serum-free differentiation (SFD) medium of IMDM supplemented with 25% Ham’s F12, 1xN2 supplement, 1xB27 supplement, 50 ug/ml ascorbic acid, 2 mM GlutaMAX, 0.4 uM monothioglycerol and 0.056% BSA at 37 °C in a 5% CO2/5% O2. hPSCs were treated with 0.05% Trypsin and plated onto low-attachment 6-well plates, and then resuspended in embryoid bodies formation medium, which is SFD medium containing 10 uM Y-27632, and 3 ng/ml human BMP4. After 12-16 h, embryoid bodies formation medium was changed into endoderm induction which is SFD medium containing 10 uM Y-27632, 0.5 ng/ml human BMP4, 2.5 ng/ml human FGF2, and 100 ng/ml human activin A for 74.5–79.5 hour. On day 4.1 or 4.3, check endoderm yield by flow cytometric analysis of CXCR4 and c-kit expression. If the endoderm yield is >90%, continue the anteriorization step. The endoderm bodies were dissociated into single cells using 0.05% trypsin and plated onto fibronectin-coated, 6-well tissue culture plates (∼7.5x10^5^ cells per well). For induction of anterior foregut endoderm, the endoderm cells were cultured in SFD medium supplemented with 100 ng/mL Noggin and 10 uM SB431542 for 24 hour, and then switched for 24 hours to 10 uM SB431542 and 1 uM IWP2 treatment in normoxic incubator. At the end of anterior foregut endoderm induction, cells were treated with ventralization media (branching media) which is SFD media containing 3 uM CHIR99021, 10 ng/ml human FGF10, 10 ng/ml human KGF, 10 ng/ml human BMP4 and 50 nM ATRA for 48 hour and three-dimensional clump formation was observed. The clumps were then suspended by gently pipetting around the wells and plated onto low-attachment plates. The suspended clumps are called lung bud organoids (LBOs) hereafter. LBOs were fed every other day until day 20–day 25 in branching media. The day 20–day 25 LBOs were embedded in 100% Matrigel in 24-well Transwell inserts and incubated in branching media in a normoxic incubator.

The directed differentiation of HIOs and HCOs were previously published^27,28^. Briefly, pluripotent stem cells were differentiated into definitive endoderm using 72 hours of Activin A (100 ng/mL) in RPMI 1640 supplemented with NEAA and increasing concentrations of defined FBS (0%, 0.2%, 2%). Spontaneous 3D mid/hindgut spheroids were generated after 96 hours of CHIR (3 uM) and FGF4 (500 ng/mL). Media was changed daily. After embedding in basement membrane Matrigel, spheroids were patterned in EGF (100 ng/mL) for HIOs or EGF and BMP2 (100 ng/mL) for HCOs for 72 hours. Both HIOs and HCOs were then maintained in Advanced DMEM/F12 supplemented with 1xN2 supplement, 1xB27 supplement, HEPES, 1X Antibiotic-Antimycotic, 1X GlutaMAX and EGF (100ng/ml) from day 10 to day 35.

#### Transplantation of human lung, intestine, and colon organoids

NSG mice were kept on antibiotic chow (275 p.p.m. Sulfamethoxazole and 1,365 p.p.m. Trimethoprim, Test Diet). Food and water were provided *ad libitum* before and after surgeries. A single conventional HIO, HCO, vHLuO, vHIO, or vHCO matured *in vitro* for 35 days, was removed from Matrigel and transplanted under the kidney capsule as previously described^14,70,71^. For the conventional HLuO transplantation, one million day 20-day 25 LBO cells were combined with 5ul of Matrigel prior to surgical implantation under the kidney capsule. Briefly, the mice were anesthetized with 2% inhaled isoflurane, and the left side of the mouse was then prepped in sterile fashion with isopropyl alcohol and povidine-iodine. A small left-posterior subcostal incision was made to expose the kidney. A subcapsular pocket was created and the organoid was then placed into the pocket. The kidney was then returned to the peritoneal cavity and the mice were given an intraperitoneal (IP) flush of Zosyn (100 mg/kg.). The skin was closed in a double layer and the mice were given a subcutaneous injection with Buprenex (0.05 mg/kg). Mesentery transplantation of vHIO or vHCO was performed as described previously^71^. Briefly, a 2 cm midline incision was made to access the abdominal cavity. The cecum was identified and removed from the body cavity with the intestine following. The mesentery was splayed out, and an appropriate location with bifurcating mesenteric vessels was identified as the transplantation site. Here, octyl/butyl cyanoacrylate adhesive glue was applied and the organoid was seeded. After allowing the glue to cure, the bowel was returned to the abdominal cavity and the incision was closed. At 8-15 weeks following engraftment, the mice were then humanely euthanized and the resulting grafts were analyzed.

#### Generation of organotypic endothelium in vessel organoid

iPSC-vessel organoids (VOs) were generated based on a protocol described by Wimmer et al.^43^ with some modifications. iPSCs were first dissociated into single cells and resuspended in Aggregation media. 1.2x10^6^ cells were added to each well of the Aggrewell™400 that were prepped with Anti-Adherence solution per manufacturer’s instructions. The Aggrewell™400 consisting of cells was then spun down at 100g for 3 mins to evenly distribute cells into each microwell (1000 cells per microwell). Cells were then left to form EBs overnight at 37°C. The next day, EBs were transferred to N2B27 media supplemented with 12μM CHIR99021 and 30ng/ml BMP4 and cultured on an orbital shaker for three days. Subsequently, the media was changed to N2B27 supplemented with 2μM Forskolin and 100ng/ml VEGFA-165 for two days. Then from days 5-15, the aggregates were further differentiated and cultured in Blood Vessel Induction (BVI) media consisting of StemPro™-34 SFM media supplemented with StemPro™-34 nutrient mix, 1X GlutaMAX, 1X Antibiotic-Antimycotic, 15% Fetal Bovine Serum (FBS), 100ng/ml VEGFA-165, and FGF2. For ligand-treated conditions, the appropriate concentrations of ligands were added to fresh BVI media and fresh BVI media was replaced every two days until the end of 15 days of differentiation. WNT2B: 50ng/ml; WNT11: 200ng/ml; SEMA3C: 200ng/ml; SEMA6B: 100ng/ml; SEMA6A: 200ng/ml; WNT4: 50ng/ml; ANGPTL4: 1ug/ml; TGFB1: 5-10ng/ml; GDF15: 10-20ng/ml.

#### EC isolation from iPSC-vessel organoids

Day 15 iPSC-VOs were pooled together and dissociated into single cells via enzymatic digestion. A cocktail of 0.5mg/ml Liberase™ TH, 5mg/ml Dispase in DPBS were utilized. The reaction was then incubated at 37°C for 20-30 mins. After the first 5 mins, 60U/ml DNase I was added to the reaction and organoids were broken down by pipetting using a wide bore tip every 5 mins thereafter. After single cells were obtained, the digestion reaction was quenched using cold KnockOut DMEM/F12 supplemented with 10% FBS. Cells were spun down at 300g for 5 mins and resuspended into MACS® buffer consisting of 0.5% MACS® BSA Stock Solution and 2mM EDTA in DPBS. The cells were passed through a 40μm filter to ensure single cells were obtained. CD144-expressing ECs were then isolated using the human CD144 (VE-Cadherin) MicroBeads and LS Columns according to manufacturer’s instructions.

#### Flow cytometry analysis

Day 3 and Day 7 spheroids/organoids were dissociated with 1X TrypLE Express Enzyme for 7∼10 mins at 37°C. The reaction was vortexed every other minute and stopped by adding N2B27 media. Single-cell suspension was made by filtering the cells through 40um Flowmi. The following antibodies were used for staining: anti-CD117 (c-KIT)-PE (1:100 dilution), anti-CD184 (CXCR4)-APC (1:100 dilution), anti-CD326 (EpCAM)-APC (1:100 dilution). For cell-surface marker staining and analyses, cells were stained for 20 min at 4°C in FACS buffer consisting of 1X PBS with 0.5% bovine serum albumin and 2mM EDTA. Stained cells were analyzed using an LSR II Flow cytometer. Data were analyzed using FlowJo software (V10.8.0).

#### Histology examination

Transplanted organoids were freshly collected, fixed in 4% PFA in 1X PBS overnight, and processed for paraffin embedding (FFPE), sectioning, H&E staining, and histological analysis. For in vitro cultured organoids, the embedded organoids were fixed in 4% PFA for 2 hours, then washed three times with 1X PBS and immersed in 30% sucrose in PBS overnight at 4°C. Afterward, the organoids were immersed in 50/50 30% sucrose/OCT for 2h at room temperature and embedded in OCT.

##### Immunofluorescence staining

For FFPE embedded sample, the sections (5um) from organoids were deparaffinized and rehydrated with a series of alcohol solutions of decreasing concentrations (5 min in each solution, 100%, 95%, 70%, and 50%) and kept in PBS for 30 min and moved to antigen retrieval process. For OCT embedded sample, the section (7um) from organoids were washed in PBS for 5 min and then fixed in 10% Neutral buffered formalin (NBF) for 10 min. Slides were washed thrice for 5 min in PBS and then moved to antigen retrieval. Antigen retrieval was performed on sections immersed in citrate 10 mM in PBS, pH 6 and microwave heated (7 min once at 650 W and 5 min twice at 350 W). Sections were washed for 10 min in PBS and then incubated for 60 min at room temperature (RT) with blocking buffer (PBS containing 4% donkey serum and 0.1% Triton X-100). Then the sections were incubated with the primary antibodies diluted in blocking buffer at 4°C overnight. After three washes (5 min) in PBS, slides were incubated for 60 min at RT with secondary antibody diluted in blocking buffer. Slides were then washed three times for 5 min in PBS and mounted in aqueous mounting medium with DAPI. The first antibodies used are listed in Key Resource Table.

##### Whole-mount immunofluorescence staining

The whole-mount staining procedure was modified following the previously published protocol^43^. Day 3 and Day 7 spheroids or organoids were collected in 15ml tube and fixed in 4% PFA at RT for one hour. The fixed samples were washed with 1X PBS and blocked with blocking buffer for 2 hours at RT (To make 50ml buffer, add 1.5ml FBS, 0.5g BSA, 250ul Triton X-100, 250ul Tween 20, and 500ul of 1% (weight/volume) sodium deoxycholate solution in 47.5ml 1X PBS). After removing the blocking buffer, 50ul of diluted first antibody solution was added overnight at 4°C. On the next day, the samples were washed three times for 15 mins with PBS-T (Tween-20, 0.05%) and incubated with fluorescent dye-conjugated secondary antibodies and DAPI at RT for two hours. After the secondary antibody reaction, the samples were washed three times with PBS-T and cleared with Benzyl Alcohol/ Benzyl Benzoate solution (BABB) for immediate imaging.

##### Immunohistochemistry staining

Sections were deparaffinized, rehydrated, and washed in PBS. Endogenous peroxidase activity was quenched with 0.3% hydrogen peroxide in methanol. Sections were preincubated with the appropriate donkey serum and then incubated with primary antibody overnight at 4 °C. The antibody used in this investigation was HOPX Rabbit pAb (1:100). The slides were treated with streptavidin-biotin complex for 60 minutes at a dilution of 1:100. Immunoreactions were visualized using a 3,3′-diaminobenzidine (DAB) substrate-chromogen solution and counterstained with eosin. Sections were immersed in an ethanol and xylene bath and mounted for examination.

#### Single molecule fluorescence in situ hybridization (smFISH)

smFISH was performed using a proprietary high-sensitivity RNA amplification and detection technology, according to the manufacturer’s instructions using the indicated proprietary probes, the RNAscope Multiplex Fluorescent Reagent Kit (v.2), and Opal dyes (1:500 dilution for Opal 570 and 690 dyes, 1:250 dilution for Opal 520). After smFISH, sections were mounted in aqueous mounting medium with DAPI. Proprietary (Advanced Cell Diagnostics) probes used were listed in Table S13.

#### Quantitative reverse-transcription PCR

Our detailed protocol was previously described^72^. Briefly, total RNA was extracted, purified, and quantified for reverse transcription using High-Capacity RNA to cDNA Kit according to the manufacturer’s instructions. qPCR was carried out using 5 ng cDNA and 6 mL SYBR green master mix. Primers were listed in Table S14. Each measurement was performed in triplicate. Heatmaps for qPCR results were generated by Morpheus visualization tool. A relative color scheme uses the minimum and maximum values in each treatment to convert values to colors. N=3 in each treatment group.

#### Proteomics mass spectrometry analysis

The fluid from two 3-month tvHLuO was collected with a 5ml syringe, combined, and immediately snap-frozen before analysis. A 1-D silver stained (SS) gel was run on the samples to check the protein levels. For protein identification, a preparative gel using 20 ug of each sample was run into a 4-12% B-T gel using MOPS buffer with molecular weight marker lanes in between each sample just until all the protein entered the gel (about 2 cm total). The full section represented the 2 cm run in the wells, were excised, reduced with DTT, alkylated with IAA, and digested with trypsin overnight. The resulting peptides were extracted and dried in a SpeedVac concentrator, then resuspended in 0.1% Formic acid (FA). 0.5ug of each sample was analyzed by nano LC-MS/MS (Orbitrap Eclipse) with data recorded using Xcalibur 4.3 software. Full details of the LC-MS methodology were reported previously^73^. The resulting spectra were searched against a combined database of common proteomics contaminant proteins (e.g., trypsin, BSA, cytokeratins), and the UniProt mouse (UP000000589) and Human (UP000005640) database using Proteome discoverer version 3.0 with the Sequest HT search algorithm.

#### Processing and analysis of 10x single-cell RNA-seq of vascularized human organoid

Day-3 meso-endoderm spheroids (B0, B1, B2, and B3) and day-7 vascularized anterior foregut (vAFG-‘B0’ and vAFG-‘B1’) and mid/hindgut organoids (vMHG-‘B1’, vMHG-‘B2’, and vMHG-‘B3’) were dissociated with 1 X TrypLE Express Enzyme at 37°C for 7–12 mins. The reactions were stopped by adding N2B27 basal medium and centrifuged and filtered with 40um strainer. Day-21 vascularized gut tube organoids (vHLPO, vHIO, and vHCO) were embedded in a Collagen-Matrigel mixture, so were dissociated with a different digestion cocktail. Briefly, the organoids from one trans-well insert were immersed in 1ml Cell Recovery Solution at 4°C for 5 mins and triturated with P1000 wide-bore tip to break the matrix. The solution was centrifuged and resuspended in 1ml pre-warmed Dispase (5mg/ml)/Liberase solution (0.5mg/ml), and the organoids were minced with scissors. The mixture was incubated at 37°C for 20 mins with gentle triturate every 5 – 7 mins. After the first 5 mins, 10ul Dnase I (60U/ml) was added to the reaction. To stop the reaction, 9ml of Knock-out DMEM/F12 containing 10% FBS was added and centrifuged and filtered with 40um strainer. Cell pellets were resuspended in N2B27 basal medium and loaded to 10x Chromium controller for 3’ single-cell RNA-seq (v3.1).

Sequencing reads from 10x Chromium scRNA-seq of individual samples were preprocessed, aligned, and quantified using 10x Genomics Cell Ranger (v6.1.2) and human reference hg38 (10x Genomics refdata-gex-GRCh38-2020-A). 10x Cell Ranger filtered data were used for downstream QC assessment and filters were then applied to keep cells with at least 500 genes, less than 100,000 unique molecular identifiers (UMIs), and less than 25% UMIs mapped to mitochondrial genes. Downstream analyses were performed in R v4.1.0 and python 3.

Analysis of individual human organoid scRNA-seq samples was performed using Seurat v4.2.0^74^. Data normalization and highly variable gene (HVG) identification were performed using SCTransform function with the effect of cell cycle scores and mitochondrial percentage regressed out. Zscore-scaled expression of HVGs was used for principal component analysis (PCA). The top 50 PCA dimensions with the largest variances were used for clustering analysis using Seurat FindClusters function and generation of UMAP visualizations using Seurat RunUMAP function. Cell clusters in each sample were identified using the Leiden algorithm^75^. Genes selectively expressed in each cluster were identified using Seurat FindAllMarkers function with two-tailed Wilcoxon rank sum test and the following criteria: fold change >=1.5, Bonferroni adjusted p value<0.1, and expression percentage >=20%.

Integrations of human organoid scRNA-seq data from different samples were performed using Harmony v0.1.0^76^, including the integration of Day 3 B0-B3 samples, the integration of Day 21 vHIO and vHCO data, and the integration of the predicted mesenchymal cells or endothelial cells (ECs) from Day 21 vHLPO, vHIO, and vHCO samples. In each of the above integrations, raw gene expression count in cells of the selected samples and cell populations were extracted and merged, normalization and HVG identification were performed using Seurat SCTransform function with the effect of cell cycle scores and mitochondrial percentage regressed out, and PCA analysis was performed using the zscore-scaled expression of HVGs. Harmony based data integration was performed on the PCA dimensions. The top 200 PCA dimensions were used in all the integrations except the integration of Day 21 ECs that used the top 50 PCA dimensions. Harmony-corrected dimensions were used for generation of UMAP visualizations using Seurat RunUMAP function and cell clustering analysis using Seurat FindClusters function using the Leiden algorithm. Genes selectively expressed in each cluster were identified using Seurat FindAllMarkers function with two-tailed Wilcoxon rank sum test and the following criteria: fold change >=1.5, Bonferroni adjusted p value <0.1, and expression percentage >=20%. For the integrated Day 21 vHIO/HCO data, a small group of cells (n=17) were separated from all other cells in the UMAP projection and selectively expressed multiple immune marker genes (e.g., *CD14*, *CD163*, *CD84*, *SPI1*), we refined the cell identity of this group of cells as immune cells.

RNA velocity analysis of the integrated Day 3 samples was performed using velocyto v0.17.16^77^ in python 3.6.3 for generation of the spliced and unspliced count matrices and using scVelo v0.2.4^78^ with steady mode in python 3.9.16 for RNA velocity estimation and visualization.

Ligand-receptor based cell-cell communication analysis of Day 21 samples was performed using CellChat v1.4.0^79^. Cells annotated with “Low-quality” were not included in the analysis. Significant interactions were inferred using data from individual samples. Interactions for a sample involving cell types with less than 5 cells in the sample were excluded. Interactions from individual samples were merged for comparison and heatmap visualization based on the inferred interaction probabilities.

#### Processing and analysis of published datasets

##### scRNA-seq of CS7 human embryo^17^

Raw gene expression count matrix, cell cluster annotation, and UMAP coordinates of CS7 human embryo data were downloaded from http://www.human-gastrula.net/. UMAP visualization was generated using the downloaded original UMAP coordinates. Prediction of CS7 cell types in the Day 3 human organoid scRNA-seq cells was performed using SingleR v1.6.1^80^.

##### scRNA-seq of E6.5-8.5 mouse embryo^18^

Raw gene expression counts, cell metadata, and PCA dimensions were downloaded as a compressed file “atlas_data.tar.gz” from https://github.com/MarioniLab/EmbryoTimecourse2018. We converted mouse symbols to human symbols based on the human and mouse homology mapping (HOM_MouseHumanSequence.rpt) downloaded from Mouse Genome Informatics (www.informatics.jax.org), used the count matrix with the converted human symbols for Seurat v4 object construction, normalized data using Seurat SCTransform function, performed supervised PCA analysis (SPCA) on the original PCA dimensions, and used the SPCA dimensions to generated a UMAP model using Seurat RunUMAP function with “return.model=T”. The constructed Seurat object with the UMAP model were used for reference mapping of Day 3 human organoid scRNA-seq using Seurat FindTransferAnchors and MapQuery functions, including transferring mouse embryo cell type labels to human organoid cells and projecting human organoid cells to the UMAP coordinates of the mouse data.

Data of mouse E6.5∼E8.5 ‘Gut’ subtypes were extracted from the above whole dataset by selecting cells with the following “endo_gutCluster” annotations, including Foregut 1, Foregut 2, Hindgut 1, Hindgut 2, Midgut, Midgut/Hindgut, Pharyngeal endoderm. Data were used to construct a Seurat v4 object with SCTransform based normalization followed by PCA analysis and UMAP model generation. The constructed Seurat object with the UMAP model were used for reference mapping of Day 3 human organoid scRNA-seq using Seurat FindTransferAnchors and MapQuery functions, including transferring mouse gut cell type labels to human organoid cells and projecting human organoid cells to the UMAP coordinates of the mouse data.

##### scRNA-seq of E8.75 mouse anterior gut data^9^

Counts matrices for cells from mouse embryos from E8.5, E9.0 and anterior and posterior E9.5 foreguts, as well as the annotated Metadata for all cells, were downloaded from NCBI GEO. Data was converted into Seurat object, and cell proportions of “LineageAnnotations” by “Stages” as defined in the Metadata were determined using Seurat’s count_table function.

##### scRNA-seq of E8.75 mouse anterior and posterior gut data^34^

Raw gene expression count matrix of all cells and cell metadata (cell index, cluster, time point, and cell type annotation) were downloaded from https://endoderm-explorer.com/. We subset the data to E8.75 anterior and posterior cells based on the original cell cluster annotation (E8.75_ap clusters) and determined anterior (n=3,579) and posterior (n=3,667) cells based on the original cell index labeling. Data normalization and dimension reduction were performed using Seurat SCTransform function followed by RunPCA function. Anterior and posterior data were integrated using Harmony v0.1.0^76^ using the top 200 PCA dimensions. UMAP visualization was generated using Seurat RunUMAP function using the harmony-corrected dimensions.

To assess the correlation of anterior and posterior cell types, we created a pseudo-bulk gene expression profile for each of the anterior and posterior cell types by averaging the expression of each gene in all cells in the given cell type, found highly variable genes (HVGs) among pseudo-bulk profiles using Seurat FindVariableFeatures function, performed PCA analysis using zscore-scaled pseudo-bulk expression of HVGs, and assessed Pearson’s correlation among cell types based on the top 10 PCA dimensions. Hierarchical clustering analysis and heatmap visualization were performed using R package pheatmap (v1.0.12) using the correlation matrix as input.

Differential gene expression between mouse anterior vs. posterior cells or between Day 7 human organoid vAFG-‘B1’ vs. vMHG-‘B3’ cells were identified using Seurat FindMarkers function using log normalization data and two-tailed Wilcoxon rank sum test with the following criteria: fold change >=1.2, p value <0.05, and expression percentage >=10%.

For pseudo-bulk correlation analysis with human organoid data, we converted mouse symbols to human symbols based on the human and mouse homology mapping (HOM_MouseHumanSequence.rpt) downloaded from Mouse Genome Informatics (www.informatics.jax.org). Pseudo-bulk correlation analysis of Day 7 human organoid scRNA-seq cell clusters with mouse anterior and posterior gut tube cell types were performed as follows. First, we created a pseudo-bulk gene expression profile for each of the mouse and human cell clusters by averaging the expression of each gene in all cells in the given cell cluster. Then Seurat’s FindVariableFeatures was used to find the top 2000 highly variable genes (HVGs) among the mouse pseudo-bulk profiles, denote HVG1, and the HVGs among the human organoid pseudo-bulk profiles, denote HVG2. We took the union of HVG1 and HVG2 and kept the genes that are present in both human and mouse data. We performed zscore scaling of the expression of HVGs among the human and mouse pseudo-bulk profiles, separately. Pearson’s correlations among all the pseudo-bulk profiles were calculated using the scaled expression of HVGs. Hierarchical clustering analysis and heatmap visualization were performed using R package pheatmap (v1.0.12) using the correlation matrix as input.

##### scRNA-seq of conventional human lung organoids^33^

Raw sequencing reads from 10x scRNA-seq of day 10 spheroids and 3-week lung progenitor organoids (LPOs) were downloaded from ArrayExpress using accession number E-MTAB-11953 (sample number ERS12457826 for day 10 spheroids; ERS12457827 and ERS12457828 for two biological replicates of 3-week LPOs). Sequencing reads of individual samples were preprocessed, aligned, and quantified using 10x Genomics Cell Ranger (v6.1.2) and human reference hg38 (10x Genomics refdata-gex-GRCh38-2020-A). 10x Cell Ranger filtered data were used for downstream QC assessment. The QC criteria reported in Hein et al.^33^ were used to perform cell pre-filtering: for day 10 spheroids, cells with 500-8,000 expressed genes, 500-50,000 unique molecular identifiers (UMIs), and <10% UMIs mapped to mitochondrial genes were included in the downstream analysis; for 3-week LPOs, cells with 500-7,000 expressed genes, 500-50,000 UMI counts, and <10% UMIs mapped to mitochondrial genes were included. Downstream analyses were performed using Seurat v4.2.0 in R v4.1.0. Data normalization and HVG identification were performed using SCTransform function with the effect of cell cycle scores and mitochondrial percentage regressed out. Two biological replicates of 3-week LPOs were integrated using Harmony v0.1.0. Cell clusters were identified using the Seurat FindClusters function with Leiden algorithm: resolution=0.8 for the LPO cell clusters and resolution=0.3 for the spheroid cell clusters. Cell clusters were assigned to cell types based on the expression of marker genes used in Hein et al.^33^. For the day 10 spheroid data, a small group of cells (n=15) were separated from all other cells in the UMAP projection and selectively expressed multiple endothelial marker genes (e.g., *KDR, CDH5, PECAM1, ESAM*). We refined the cell identity of this group of cells as endothelium progenitors.

##### scRNA-seq of human intestinal organoids^29^

Raw sequencing reads from 10x scRNA-seq of hindgut (‘day 0’ sample) and human intestinal organoids (HIOs, ‘day 14’ sample) were downloaded from ArrayExpress using accession number E-MTAB-11953 (sample number ERS4758594 and ERS4758595 for the ‘day 0’ and ‘day 14’ HIO data, respectively). Sequencing reads of individual samples were preprocessed, aligned, and quantified using 10x Genomics Cell Ranger (v6.1.2) and human reference hg38 (10x Genomics refdata-gex-GRCh38-2020-A). 10x Cell Ranger filtered data were used for downstream QC assessment. The following QC criteria used in the GitHub code of Holloway et al.^29^ were applied to select cells for downstream analysis, including 1,000-9,000 expressed genes, <60,000 unique molecular identifiers (UMIs), and <10% UMIs mapped to mitochondrial genes. Downstream analyses were performed using Seurat v4.2.0 in R v4.1.0. Data normalization and HVG identification were performed using SCTransform function with the effect of cell cycle scores and mitochondrial percentage regressed out. Cell clusters were identified in hindgut and HIO data, separately, using the Seurat FindClusters function using Leiden algorithm with resolution=0.8. Cell clusters were assigned to cell types based on the expression of marker genes used in Holloway et al.^29^

##### scRNA-seq of fetal human lung^30^

Gene expression matrix, cell metadata, PCA dimensions, and UMAP coordinates of all scRNA-seq of fetal human lung cells were downloaded as an h5ad file “Assembled10DomainsFiltered.h5ad” from https://fetal-lung.cellgeni.sanger.ac.uk/. Prediction of fetal human lung cell types in Day 21 vHLPO cells was performed using SingleR v1.6.1^80^. We first performed the prediction using the “broad_celltype” annotation of fetal lung cells. Predictions were grouped into four cell lineages as follows for comparison with the cluster based Day 21 vHLPO cell types: endothelial (Lymph endothelial and Vas endothelial), epithelial (Distal/Proximal epithelial), immune (T & ILC, B, NK, Other myeloid, Meg-ery), Mesen&PNS (Fibroblast, Mesothelial, Myofibro & SMC, Chondrocyte, PNS). We then iterated into the prediction of fetal lung cell types within each lineage for the Day 21 vHLPO cell clusters.

##### scRNA-seq of developing human endodermal organs^35^

scRNA-seq count matrix and metadata of fetal atlas cells (n=155,232) were downloaded from Mendeley data using DOI listed in the key resource table. The original “cell_type” annotation was used in the downstream analysis except that the endothelial cells (ECs) were further classified into lymphatic ECs and vascular ECs as described in the following.

We extracted gene count matrix of the predicted ECs (n=5,375) based on the “Major_cell_type” annotation, constructed a Seurat v4 object, normalized gene expression using the Seurat SCTransform function followed by a PCA analysis, integrated data from different samples using Harmony, and identified cell clusters using the Leiden algorithm. Cells (n=1,167) in the clusters with co-expression of pan-EC (*CDH5*, *CLDN5*) and immune, epithelial or fibroblast cell markers were annotated as “Contaminated” cells. Lymphatic ECs (n=279 cells) were identified based on the cell cluster with selective co-expression of marker genes: *CDH5*, *CLDN5, PROX1*, *LYVE1*, and *PDPN*. Vascular ECs (n=3,929 cells) were identified based clusters with selective co-expression of *CDH5*, *CLDN5*, *PLVAP*, and *EMCN*.

To assess the tissue identity of day 21 human organoid mesenchymal cells, we performed both a correlation analysis and a SingleR based tissue identity prediction described in the following. For the correlation analysis, we first created a pseudo-bulk gene expression profile for the mesenchymal cells in each of the fetal and organoid samples by averaging the expression of each gene in all the predicted mesenchymal cells of the given sample. For the fetal tissue data, mesenchymal cells were identified based on the original “Major_cell_type” annotation and samples of ages earlier than 80 post-conception days were used. For the organoid samples, mesenchymal cells included fibroblasts, proliferative fibroblasts, pericytes, and smooth muscle cells. Seurat’s FindVariableFeatures was used to find highly variable genes (HVGs) among the fetal tissue pseudo-bulk profiles, denote HVG1, and the HVGs among the human organoid pseudo-bulk profiles, denote HVG2. We took the intersection of HVG1 and HVG2 and kept the genes that are present in both fetal tissue and organoid data. We performed zscore scaling of the expression of HVGs among the fetal tissue and organoid pseudo-bulk profiles, separately. Pearson’s correlations among all the pseudo-bulk profiles were calculated using the scaled expression of HVGs. Hierarchical clustering analysis and heatmap visualization were performed using R package pheatmap (v1.0.12) using the correlation matrix as input.

For the SingleR based prediction of fetal tissue identity in the mesenchymal cells of day 21 organoids, we constructed a reference data object using gene expression in the predicted mesenchymal cells from ileum, jejunum, and distal lung tissues from samples at the age of 80 post-conception days. Data from different samples were integrated using Harmony for a UMAP visualization. The prediction of fetal SI or lung tissue identities for the day 21 organoid mesenchymal cells was performed using SingleR v1.6.1^80^. Using the same SingleR based approach, we also performed the prediction of fetal SI or lung tissue identities for the ECs in the day 21 human organoid samples.

Ligand-receptor based cell-cell communication analysis was performed using CellChat v1.4.0^79^ on data from the following tissues: Colon, Lung-distal, Lung-airway, Jejunum, Ileum, Duodenum. Data from the age of 80 post-conception days were used as all the above tissues have >1000 cells from the samples of this age. Original cell type annotation was used except that ECs were classified to vascular EC, lymphatic EC, and contaminant based on the analysis as described above. Mesenchyme subtypes (n=5) were merged as “Mesenchyme”. Similarly, sub-types of “Macrophage/monocyte” and “T cell/NK cell” were merged, respectively. Cells annotated with “Contaminated” or “Undefined” were not included in the analysis. Significant interactions were inferred using data from individual tissues. Interactions for a sample involving cell types with less than 5 cells in the sample were excluded. For lung tissues, interactions involving “Enteroendocrine”, “Gastrointestinal epithelium”, and “Intestinal epithelium” were excluded. Interactions from individual tissues were merged for comparison and heatmap visualization based on the inferred interaction probabilities.

##### Human cell atlases of fetal gene expression^4^

scRNA-seq gene expression count matrix and cell metadata were downloaded from GEO using accession number GSE156793. We extracted data of vascular ECs in different tissues based on the original “Main_cluster_name” annotation and identified lung vascular EC specific markers using the following approach. First, we identified genes differentially expressed in vascular EC cells within each data sample (determined based on the original “organ” and “development day” annotations) using Seurat FindMarkers with two-tailed Wilcoxon rank sum test comparing gene expression in ECs with gene expression in all other cells in the tissue. Log normalized data were used for the DE test and the following DE criteria were used: p value <0.05, fold change >=1.2 in EC, and expression percentage >=20% in EC. We took the union of EC DEGs from all tested tissues and identified lung vascular EC specific genes using the following criteria: (i) differentially expressed (Bonferroni-adjusted p value <0.1 and fold change >=1.5) in lung vascular ECs when compared to vascular ECs in all other tissues; (ii) expression percentage >=20% in lung vascular ECs; and (iii) average expression in the lung vascular EC is at least 1.25 fold higher than its average expression in vascular ECs of any of other tissues.

##### Tabula Sapiens scRNA-seq atlas^5^

Tabula Sapiens scRNA-seq data with cell metadata from 25 organs of eight normal human subjects in Tabula Sapiens were downloaded as h5ad files from figshare.com listed in the key resource table. EC cells within each tissue were determined based on the “compartment” annotation (i.e., compartment = endothelial) from the original publication^5^. We integrated ECs from all tissues using Harmony and identified cell clusters using Leiden algorithm. Cell clusters with co-expression of lymphatic markers (*CCL21*, *PROX1*, *LYVE1*, and *PDPN*) were annotated as lymphatic ECs, and the remaining cells were annotated as vascular ECs. Lung vascular EC specific markers were identified using the same approach as described in the above “Human cell atlases of fetal gene expression” section.

### QUANTIFICATION AND STATISTICAL ANALYSIS

First, normal distribution from each group was confirmed using c2 test before any comparison between groups. All data represent mean ± SEM. Statistical significance was determined by parametric tests (all data possesses equal variance p>0.05): unpaired 2-tailed t-test, ANOVA (>2 groups) with post hoc tests as indicated; or non-parametric test: Mann-Whitney (2 groups). A p value of <0.05 was considered significant. Statistical details can be found in the figure legends. For all image quantification, ImageJ software was used and n represents different specimens from different patients or control subjects. All violin plot data were tested by Bonferroni correction test and FDR<0.05 was regarded as significant. Analyses were carried out using GraphPad Prism 8.0. As for all the experiments, at least three independent experiments were performed to reach the minimal requirement for statistical significance, unless otherwise specified. Blinding or randomization was not performed unless otherwise specified. Exclusion is not applied in this study.

## Key resources table

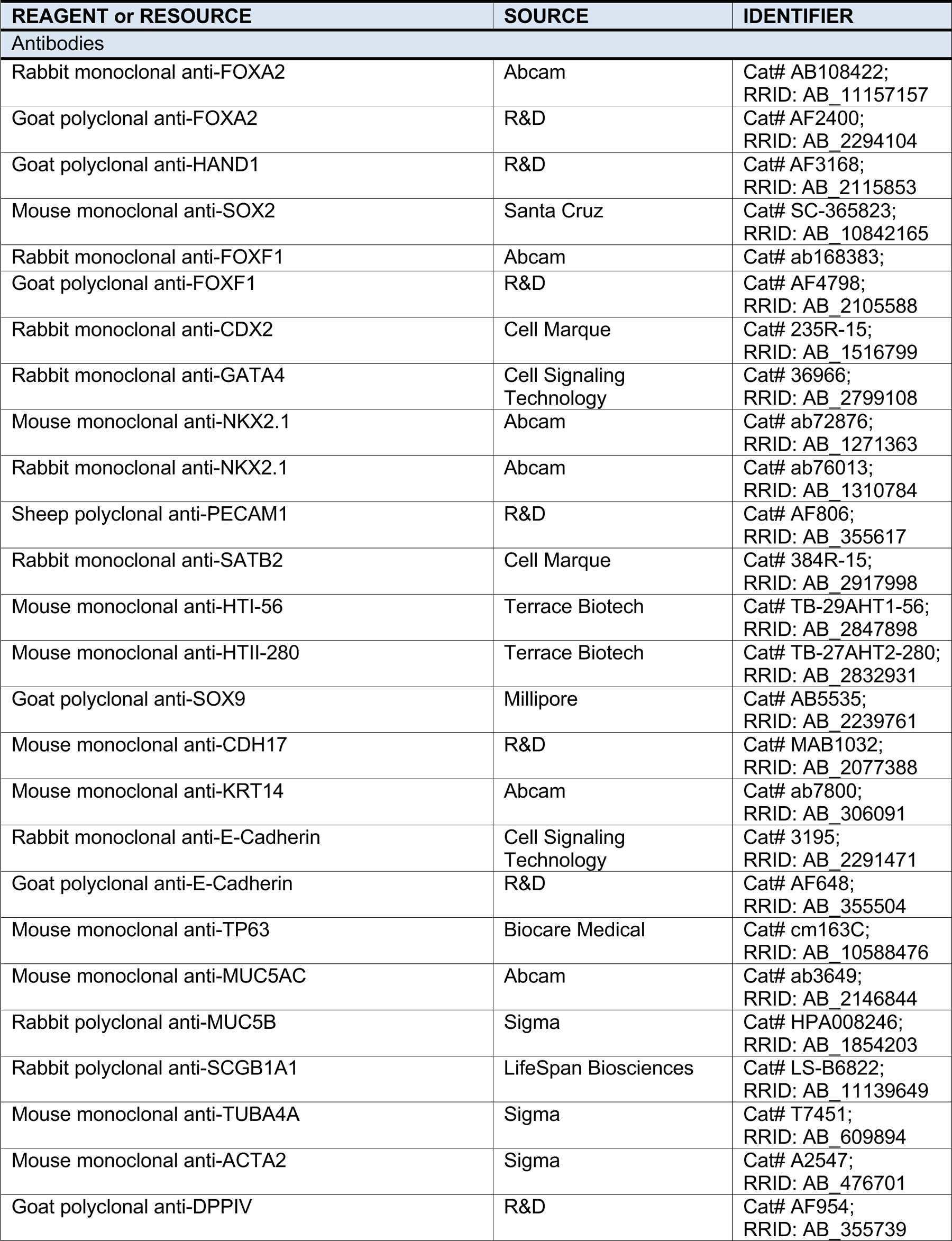
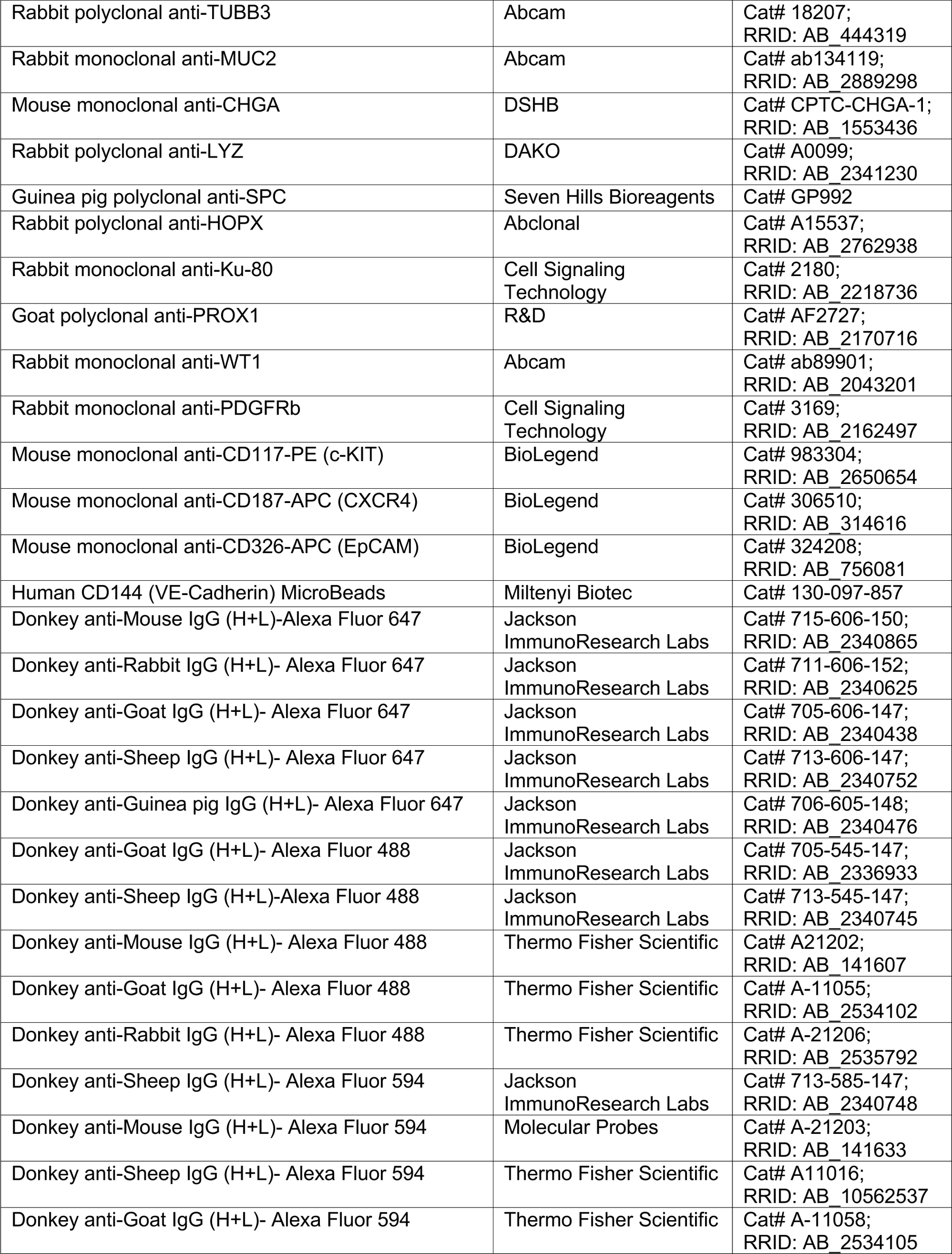
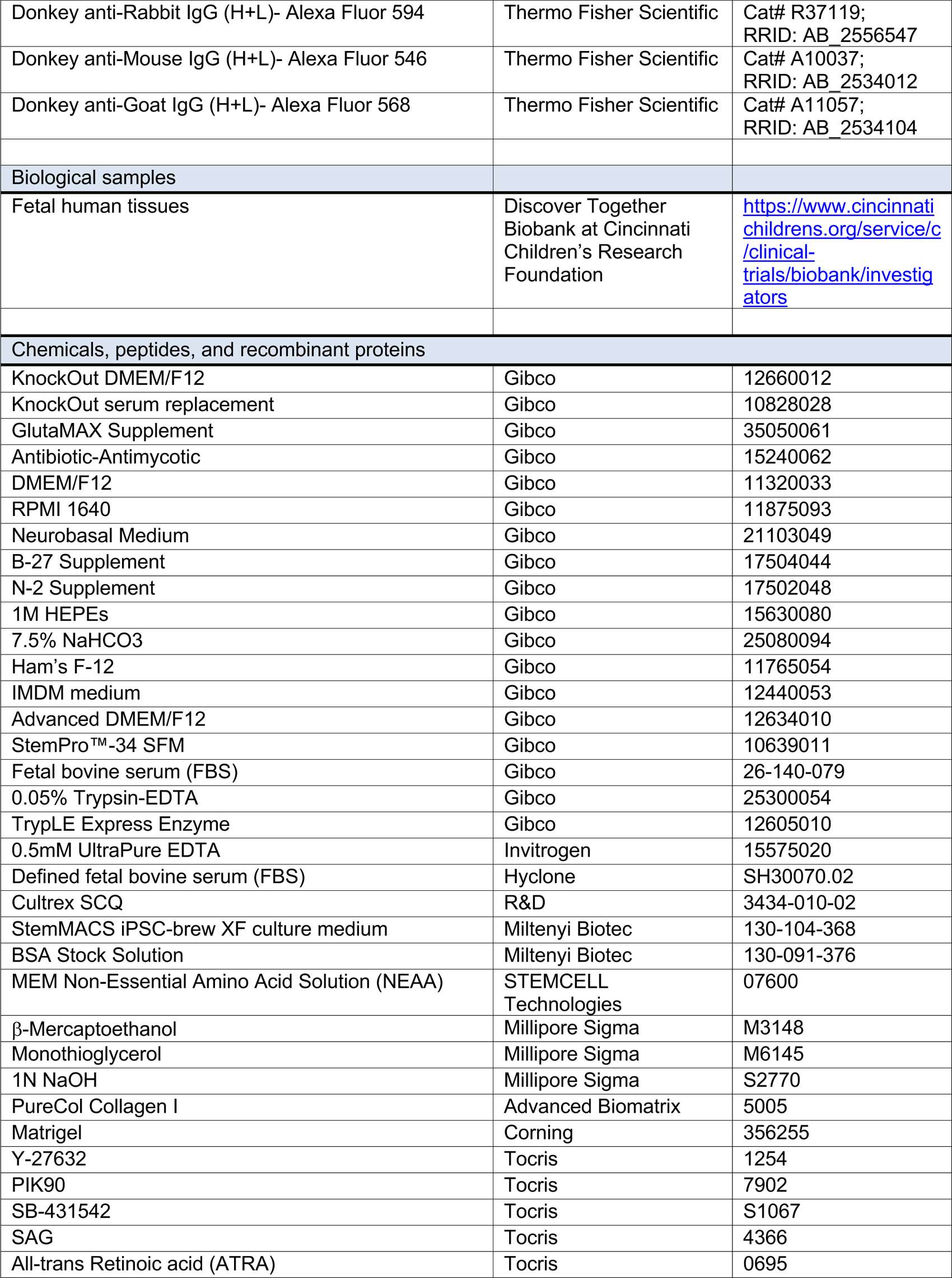
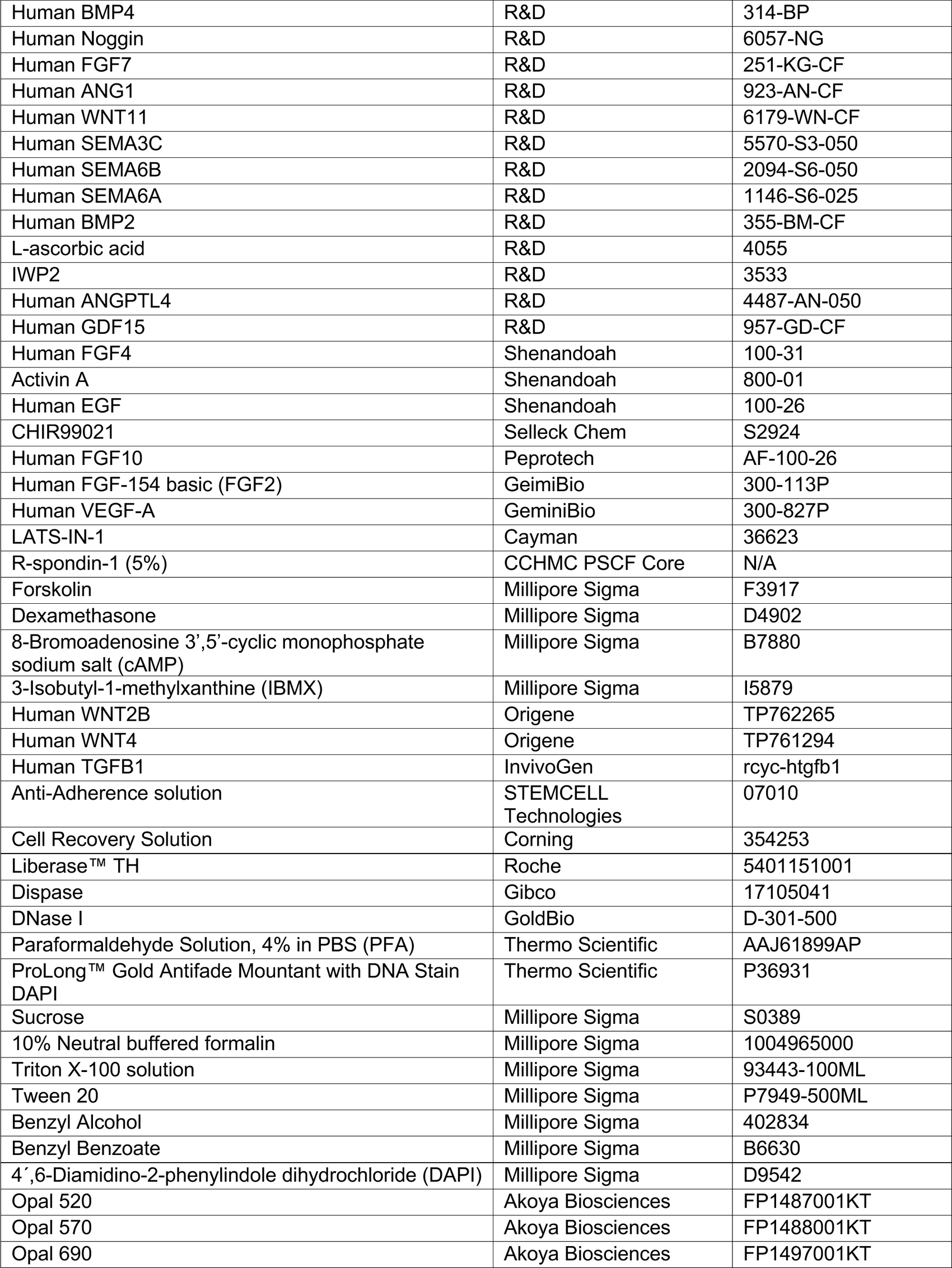
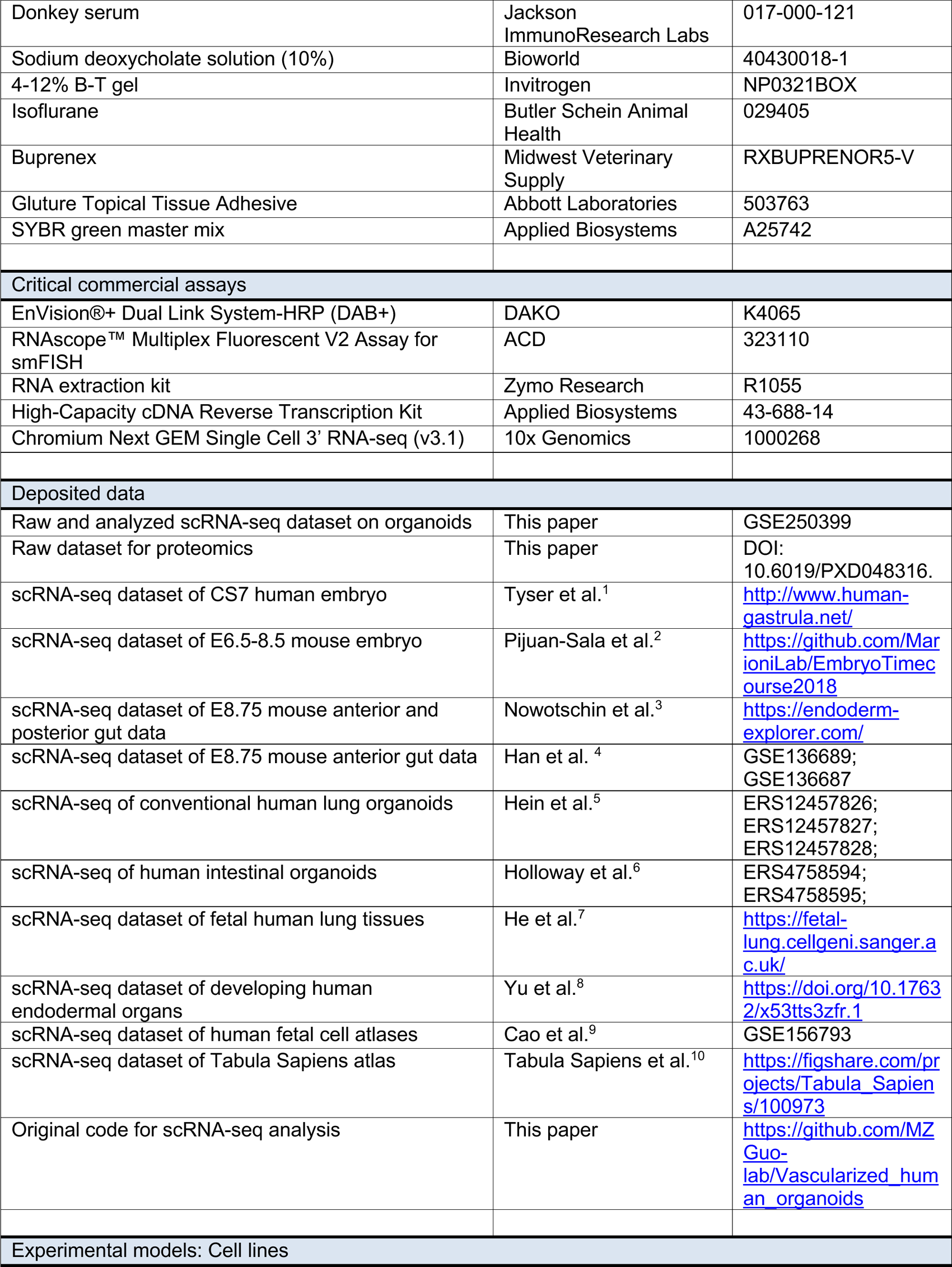
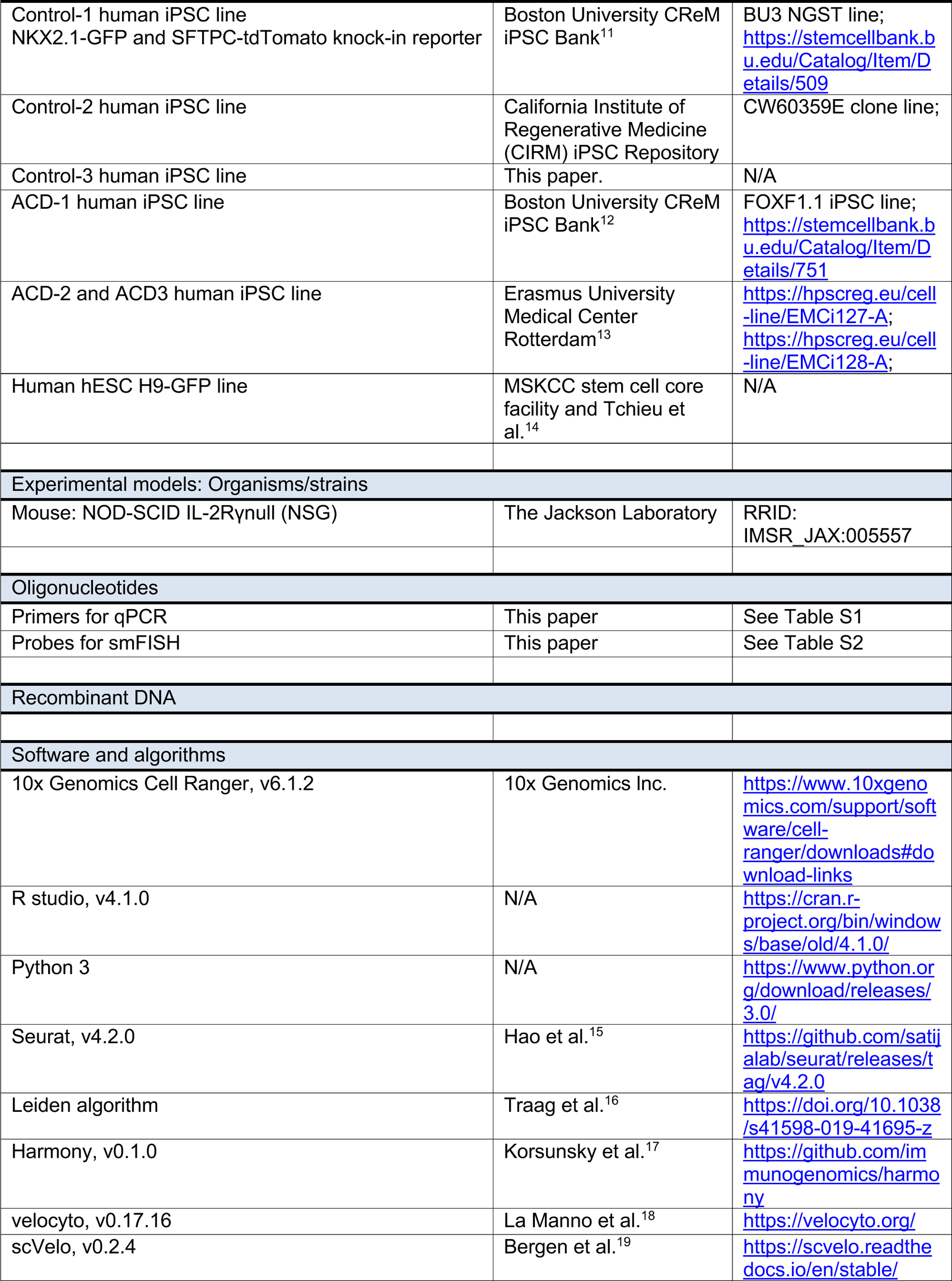
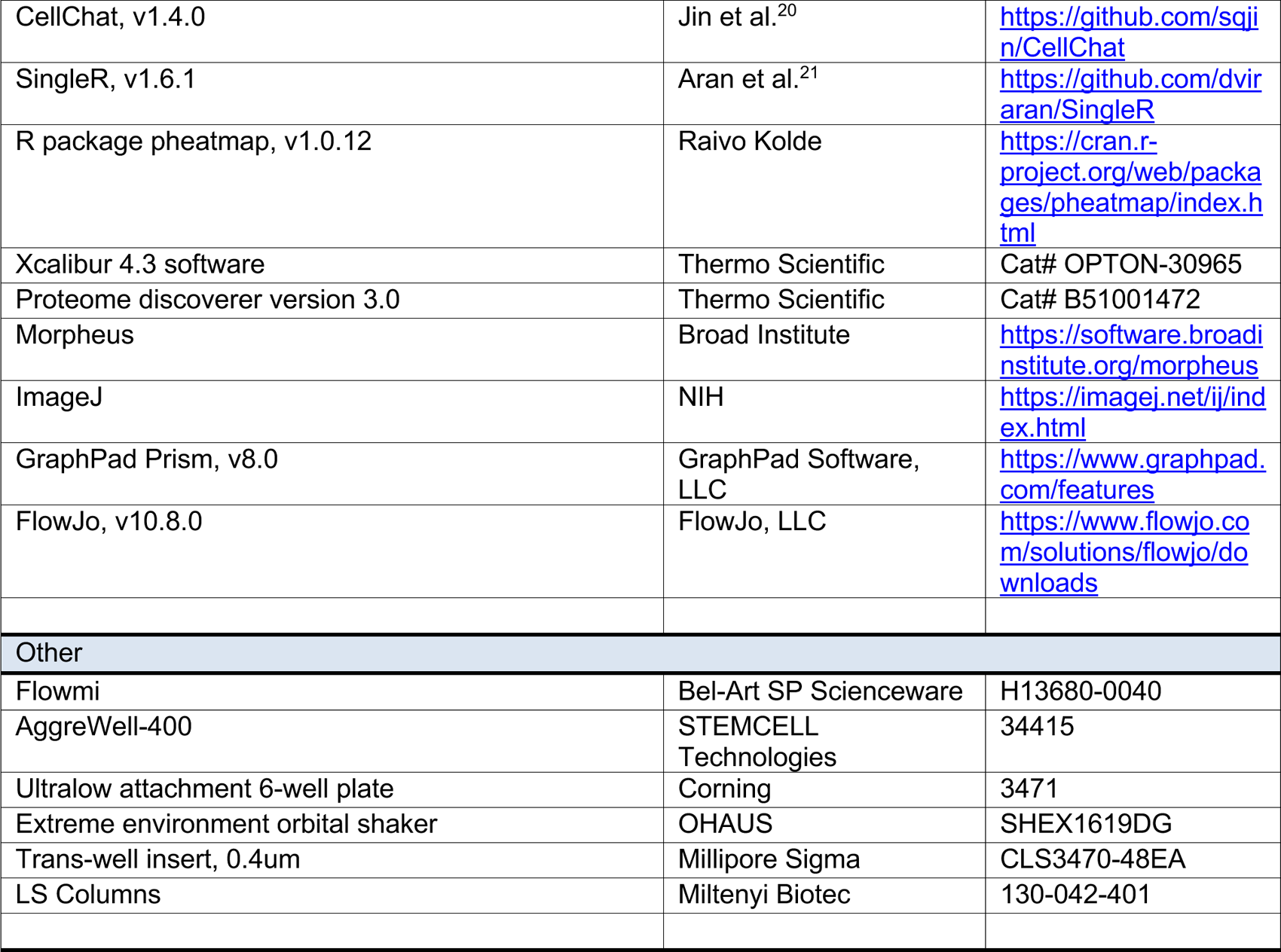

