## Supplemental Figure for "Deciphering Endothelial and Mesenchymal Organ Specification in Vascularized Lung and Intestinal Organoids"

### Figure S1

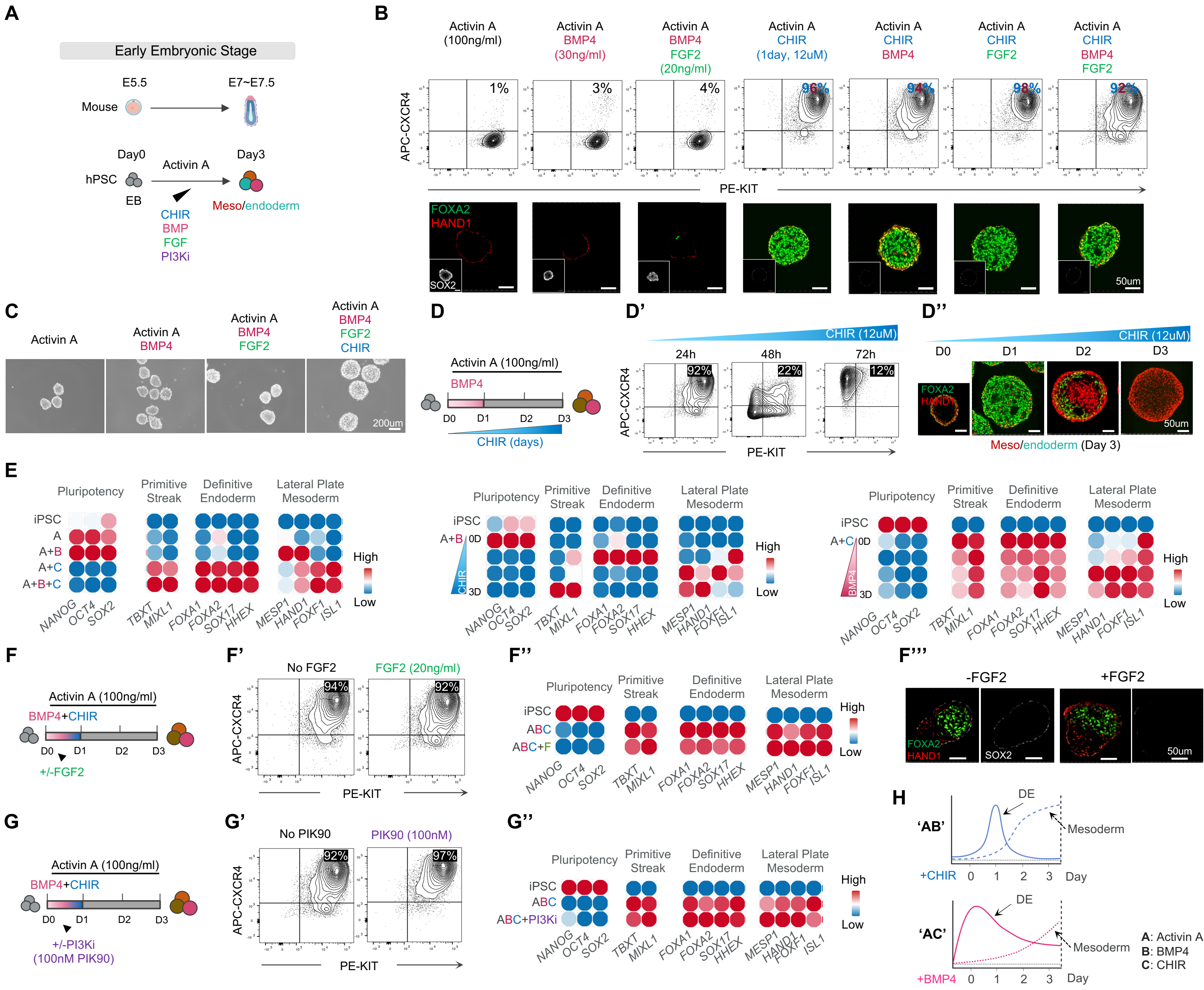
A
Early Embryonic Stage
E5.5
E7~E7.5
Mouse
Day0
Day3
Activin A
hPSC
EB
CHIR
BMP
FGF
PI3Ki
Meso/endoderm
B
Activin A (100ng/ml)
Activin A BMP4 (30ng/ml)
Activin A BMP4 FGF2 (20ng/ml)
Activin A CHIR (1day, 12uM)
Activin A CHIR BMP4
Activin A CHIR FGF2
Activin A CHIR BMP4 FGF2
APC-CXCR4
1%
3%
4%
96%
94%
98%
92%
PE-KIT
FOXA2
HAND1
SOX2
50um
C
Activin A
Activin A BMP4
Activin A BMP4 FGF2
Activin A BMP4 FGF2 CHIR
200um
D
Activin A (100ng/ml)
BMP4
D0
D1
D2
D3
CHIR (days)
D'
CHIR (12uM)
24h
48h
72h
92%
22%
12%
APC-CXCR4
PE-KIT
D''
CHIR (12uM)
D0
D1
D2
D3
FOXA2
HAND1
50um
Meso/endoderm (Day 3)
E
Pluripotency
Primitive Streak
Definitive Endoderm
Lateral Plate Mesoderm
iPSC
A
A+B
A+C
A+B+C
High
Low
NANOG OCT4 SOX2
TBXT MIXL1
FOXA1 FOXA2 SOX17 HHEX
MESP1 HAND1 FOXF1 ISL1
A+B 0D
CHIR
3D
A+C 0D
BMP4
3D
F
Activin A (100ng/ml)
BMP4+CHIR
D0
D1
D2
D3
+/-FGF2
F'
No FGF2
FGF2 (20ng/ml)
94%
92%
APC-CXCR4
PE-KIT
F''
Pluripotency
Primitive Streak
Definitive Endoderm
Lateral Plate Mesoderm
iPSC
ABC
ABC+F
High
Low
F'''
-FGF2
+FGF2
FOXA2
HAND1
SOX2
50um
G
Activin A (100ng/ml)
BMP4+CHIR
D0
D1
D2
D3
+/-PI3Ki
(100nM PIK90)
G'
No PIK90
PIK90 (100nM)
92%
97%
APC-CXCR4
PE-KIT
G''
Pluripotency
Primitive Streak
Definitive Endoderm
Lateral Plate Mesoderm
iPSC
ABC
ABC+PI3Ki
High
Low
H
'AB'
DE
Mesoderm
+CHIR
0 1 2 3
Day
'AC'
DE
Mesoderm
+BMP4
0 1 2 3
Day
A: Activin A
B: BMP4
C: CHIR

#### Figure S2

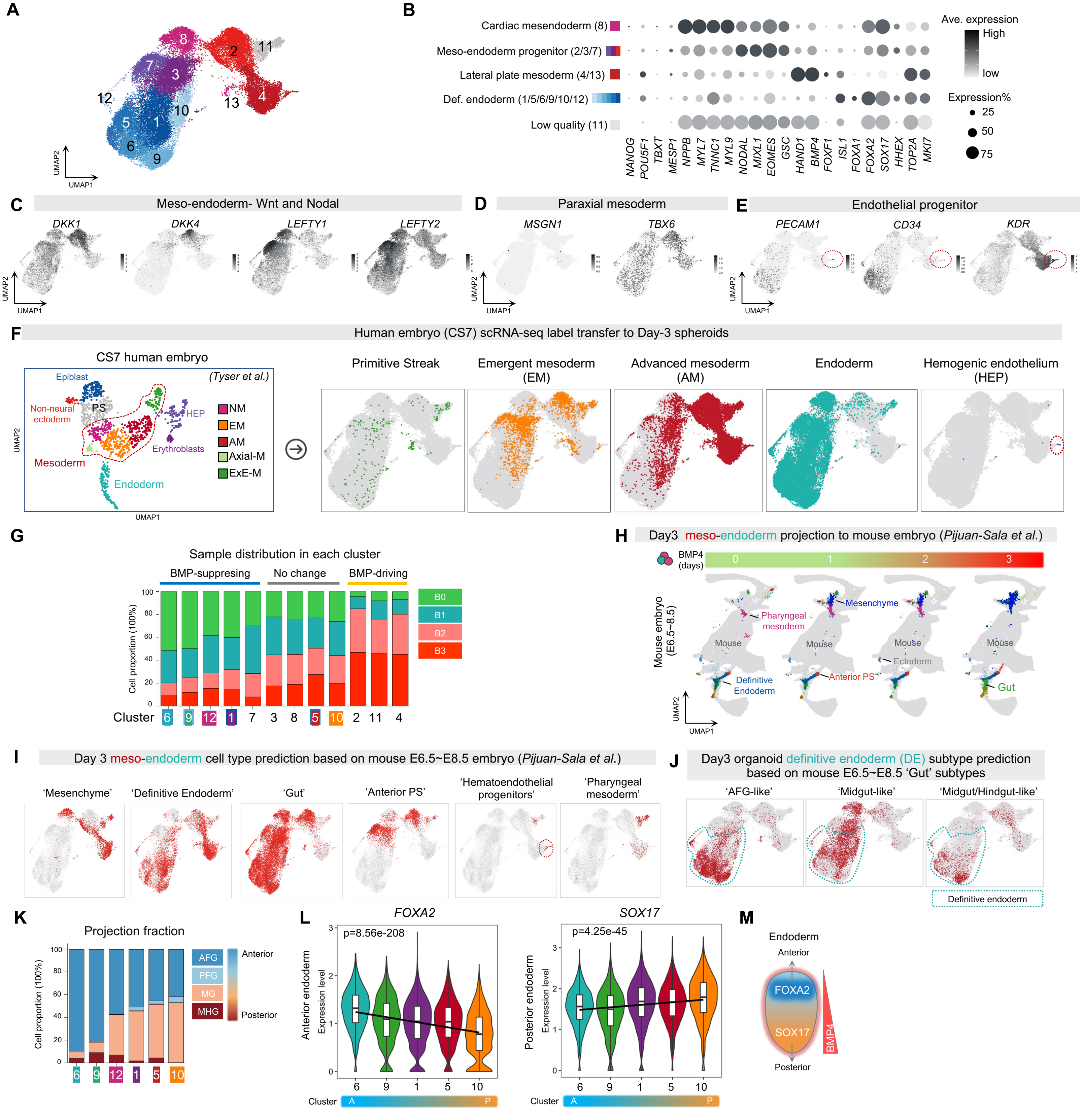

A
B
Cardiac mesendoderm (8)
Meso-endoderm progenitor (2/3/7)
Lateral plate mesoderm (4/13)
Def. endoderm (1/5/6/9/10/12)
Low quality (11)
Ave. expression
High
low
Expression%
25
50
75
C
Meso-endoderm- Wnt and Nodal
D
Paraxial mesoderm
E
Endothelial progenitor
F
Human embryo (CS7) scRNA-seq label transfer to Day-3 spheroids
CS7 human embryo
(Tyser et al.)
Primitive Streak
Emergent mesoderm (EM)
Advanced mesoderm (AM)
Endoderm
Hemogenic endothelium (HEP)
G
Sample distribution in each cluster
BMP-suppresing
No change
BMP-driving
Cell proportion (100%)
Cluster
H
Day3 meso-endoderm projection to mouse embryo (Pijuan-Sala et al.)
Mouse embryo (E6.5~8.5)
I
Day 3 meso-endoderm cell type prediction based on mouse E6.5~E8.5 embryo (Pijuan-Sala et al.)
'Mesenchyme'
'Definitive Endoderm'
'Gut'
'Anterior PS'
'Hematoendothelial progenitors'
'Pharyngeal mesoderm'
J
Day3 organoid definitive endoderm (DE) subtype prediction based on mouse E6.5~E8.5 'Gut' subtypes
'AFG-like'
'Midgut-like'
'Midgut/Hindgut-like'
Definitive endoderm
K
Projection fraction
L
FOXA2
SOX17
p=8.56e-208
p=4.25e-45
Anterior endoderm
Posterior endoderm
Expression level
M
Endoderm
Anterior
Posterior
FOXA2
SOX17
BMP4

**Figure S3**

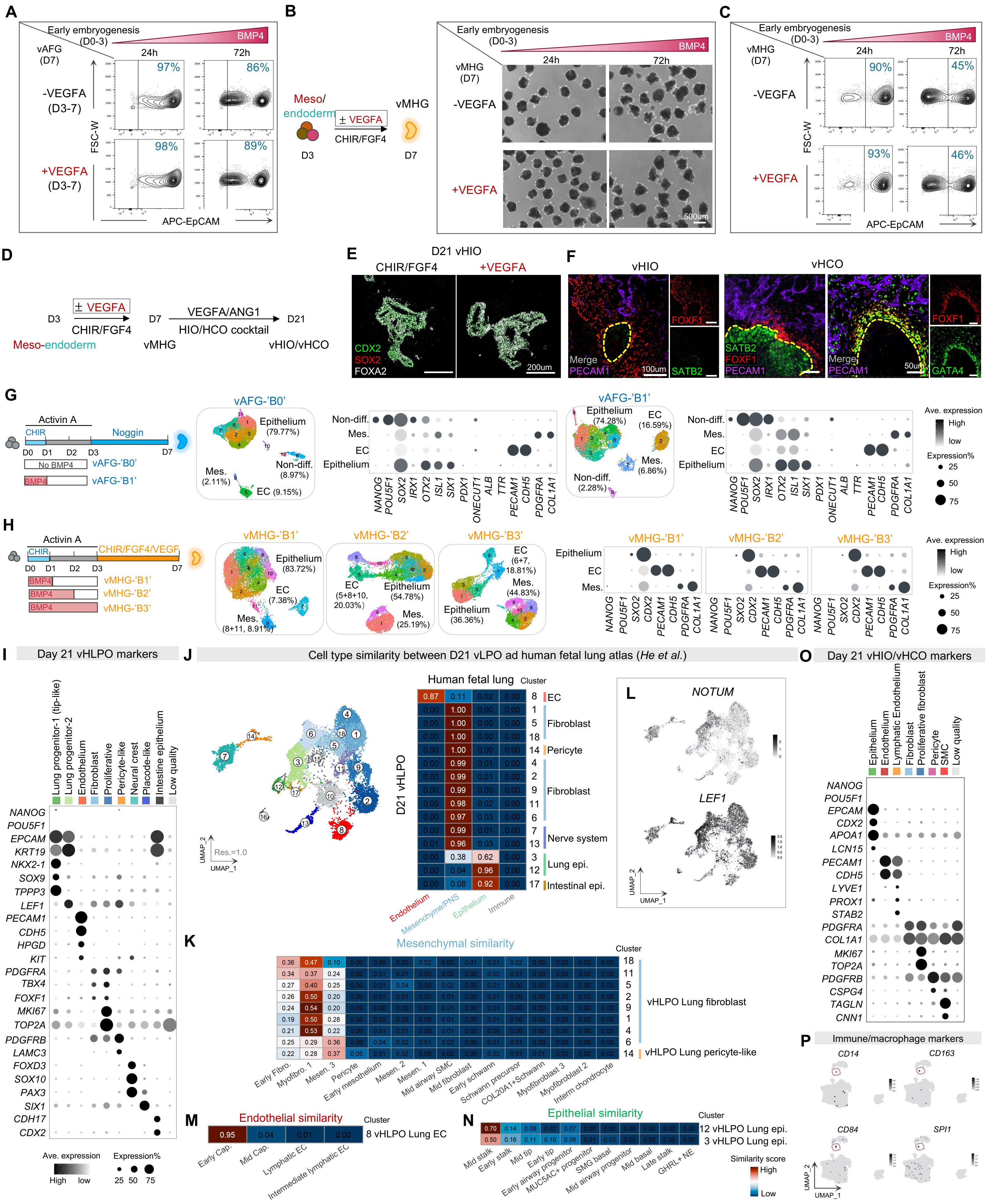
A
Early embryogenesis (D0-3)
BMP4
vAFG (D7)
24h
72h
-VEGFA (D3-7)
+VEGFA (D3-7)
97%
86%
98%
89%
FSC-W
APC-EpCAM
B
Meso/ endoderm
D3
± VEGFA
CHIR/FGF4
vMHG
D7
Early embryogenesis (D0-3)
BMP4
vMHG (D7)
24h
72h
-VEGFA
+VEGFA
500um
C
Early embryogenesis (D0-3)
BMP4
vMHG (D7)
24h
72h
-VEGFA
+VEGFA
90%
45%
93%
46%
FSC-W
APC-EpCAM
D
± VEGFA
D3
CHIR/FGF4
D7
VEGFA/ANG1
HIO/HCO cocktail
D21
Meso-endoderm
vMHG
vHIO/vHCO
E
D21 vHIO
CHIR/FGF4
+VEGFA
CDX2
SOX2
FOXA2
200um
F
vHIO
vHCO
Merge
PECAM1
FOXF1
SATB2
100um
SATB2
FOXF1
PECAM1
Merge
PECAM1
50um
GATA4
G
Activin A
CHIR
Noggin
D0 D1 D2 D3 D7
No BMP4
vAFG-'B0'
BMP4
vAFG-'B1'
vAFG-'B0'
Epithelium (79.77%)
Mes. (2.11%)
Non-diff. (8.97%)
EC (9.15%)
Non-diff.
Mes.
EC
Epithelium
NANOG POU5F1 SOX2 IRX1 OTX2 ISL1 SIX1 PDX1 ONECUT1 ALB TTR PECAM1 CDH5 PDGFRA COL1A1
vAFG-'B1'
Epithelium (74.28%)
EC (16.59%)
Mes. (6.86%)
Non-diff. (2.28%)
Non-diff.
Mes.
EC
Epithelium
NANOG POU5F1 SOX2 IRX1 OTX2 ISL1 SIX1 PDX1 ONECUT1 ALB TTR PECAM1 CDH5 PDGFRA COL1A1
Ave. expression
High
low
Expression%
25
50
75
H
Activin A
CHIR
CHIR/FGF4/VEGF
D0 D1 D2 D3 D7
BMP4
vMHG-'B1'
BMP4
vMHG-'B2'
BMP4
vMHG-'B3'
vMHG-'B1'
Epithelium (83.72%)
EC (7.38%)
Mes. (8+11, 8.91%)
vMHG-'B2'
EC (5+8+10, 20.03%)
Epithelium (54.78%)
Mes. (25.19%)
vMHG-'B3'
EC (6+7, 18.81%)
Mes. (44.83%)
Epithelium (36.36%)
Epithelium
EC
Mes.
vMHG-'B1'
vMHG-'B2'
vMHG-'B3'
NANOG POU5F1 SXO2 CDX2 PECAM1 CDH5 PDGFRA COL1A1
NANOG POU5F1 SXO2 CDX2 PECAM1 CDH5 PDGFRA COL1A1
NANOG POU5F1 SXO2 CDX2 PECAM1 CDH5 PDGFRA COL1A1
Ave. expression
High
low
Expression%
25
50
75
I
Day 21 vHLPO markers
Lung progenitor-1 (tip-like)
Lung progenitor-2
Endothelium
Fibroblast
Proliferative
Pericyte-like
Neural crest
Placode-like
Intestine epithelium
Low quality
NANOG
POU5F1
EPCAM
KRT19
NKX2-1
SOX9
TPPP3
LEF1
PECAM1
CDH5
HPGD
KIT
PDGFRA
TBX4
FOXF1
MKI67
TOP2A
PDGFRB
LAMC3
FOXD3
SOX10
PAX3
SIX1
CDH17
CDX2
Ave. expression
High
low
Expression%
25 50 75
J
Cell type similarity between D21 vLPO ad human fetal lung atlas (He et al.)
Res.=1.0
UMAP_2
UMAP_1
Human fetal lung
Cluster
D21 vHLPO
0.87 0.11 0.02 0.00 | 8 EC
0.00 1.00 0.00 0.00 | 1
0.00 1.00 0.00 0.00 | 5 Fibroblast
0.00 1.00 0.00 0.00 | 18
0.00 1.00 0.00 0.00 | 14 Pericyte
0.00 0.99 0.01 0.00 | 4
0.00 0.99 0.01 0.00 | 2
0.00 0.99 0.00 0.00 | 9 Fibroblast
0.00 0.98 0.02 0.00 | 11
0.00 0.97 0.03 0.00 | 6
0.00 0.99 0.01 0.00 | 7 Nerve system
0.01 0.96 0.03 0.00 | 13
0.00 0.38 0.62 0.00 | 3 Lung epi.
0.00 0.04 0.96 0.00 | 12
0.00 0.08 0.92 0.00 | 17 Intestinal epi.
Endothelium
Mesenchyme/PNS
Epithelium
Immune
L
NOTUM
LEF1
UMAP_2
UMAP_1
K
Mesenchymal similarity
Cluster
0.36 0.47 0.10 0.00 0.00 0.00 0.02 0.00 0.01 0.01 0.02 0.00 0.00 0.00 0.00 | 18
0.34 0.37 0.24 0.00 0.01 0.01 0.00 0.00 0.00 0.00 0.01 0.00 0.00 0.00 0.00 | 11
0.27 0.40 0.25 0.00 0.01 0.04 0.00 0.02 0.00 0.01 0.00 0.00 0.00 0.00 0.00 | 5
0.26 0.50 0.20 0.00 0.01 0.01 0.00 0.01 0.00 0.01 0.00 0.00 0.00 0.00 0.00 | 2
0.24 0.54 0.20 0.00 0.00 0.00 0.01 0.00 0.00 0.00 0.00 0.00 0.00 0.00 0.00 | 9
0.19 0.50 0.28 0.00 0.00 0.00 0.00 0.01 0.00 0.01 0.00 0.00 0.00 0.00 0.00 | 1
0.21 0.53 0.22 0.00 0.01 0.01 0.00 0.01 0.00 0.00 0.00 0.00 0.00 0.00 0.00 | 4
0.25 0.29 0.36 0.00 0.04 0.02 0.01 0.02 0.00 0.00 0.01 0.01 0.00 0.00 0.00 | 6
0.22 0.28 0.37 0.05 0.01 0.02 0.00 0.00 0.00 0.02 0.01 0.01 0.00 0.00 0.00 | 14
vHLPO Lung fibroblast
vHLPO Lung pericyte-like
Early Fibro.
Myofibro. 1
Mesen. 3
Pericyte
Early mesothelium
Mesen. 2
Mesen. 1
Mid airway SMC
Mid fibroblast
Early schwann
Schwann precursor
COL20A1+Schwann
Myofibroblast 3
Myofibroblast 2
Interm chondrocyte
M
Endothelial similarity
Cluster
0.95 0.04 0.01 0.00 | 8 vHLPO Lung EC
Early Cap.
Mid Cap.
Lymphatic EC
Intermediate lymphatic EC
N
Epithelial similarity
Cluster
0.70 0.14 0.08 0.02 0.07 0.00 0.00 0.00 0.00 0.00 0.00 | 12 vHLPO Lung epi.
0.50 0.16 0.11 0.10 0.09 0.01 0.03 0.00 0.00 0.00 0.00 | 3 vHLPO Lung epi.
Mid stalk
Early stalk
Mid tip
Early tip
Early airway progenitor
MUC5AC+ progenitor
SMG basal
Mid airway progenitor
Mid basal
Late stalk
GHRL+ NE
Similarity score
High
Low
O
Day 21 vHIO/vHCO markers
Epithelium
Endothelium
Lymphatic Endothelium
Fibroblast
Proliferative fibroblast
Pericyte
SMC
Low quality
NANOG
POU5F1
EPCAM
CDX2
APOA1
LCN15
PECAM1
CDH5
LYVE1
PROX1
STAB2
PDGFRA
COL1A1
MKI67
TOP2A
PDGFRB
CSPG4
TAGLN
CNN1
P
Immune/macrophage markers
CD14
CD163
CD84
SPI1
UMAP_2
UMAP_1

#### Figure S4

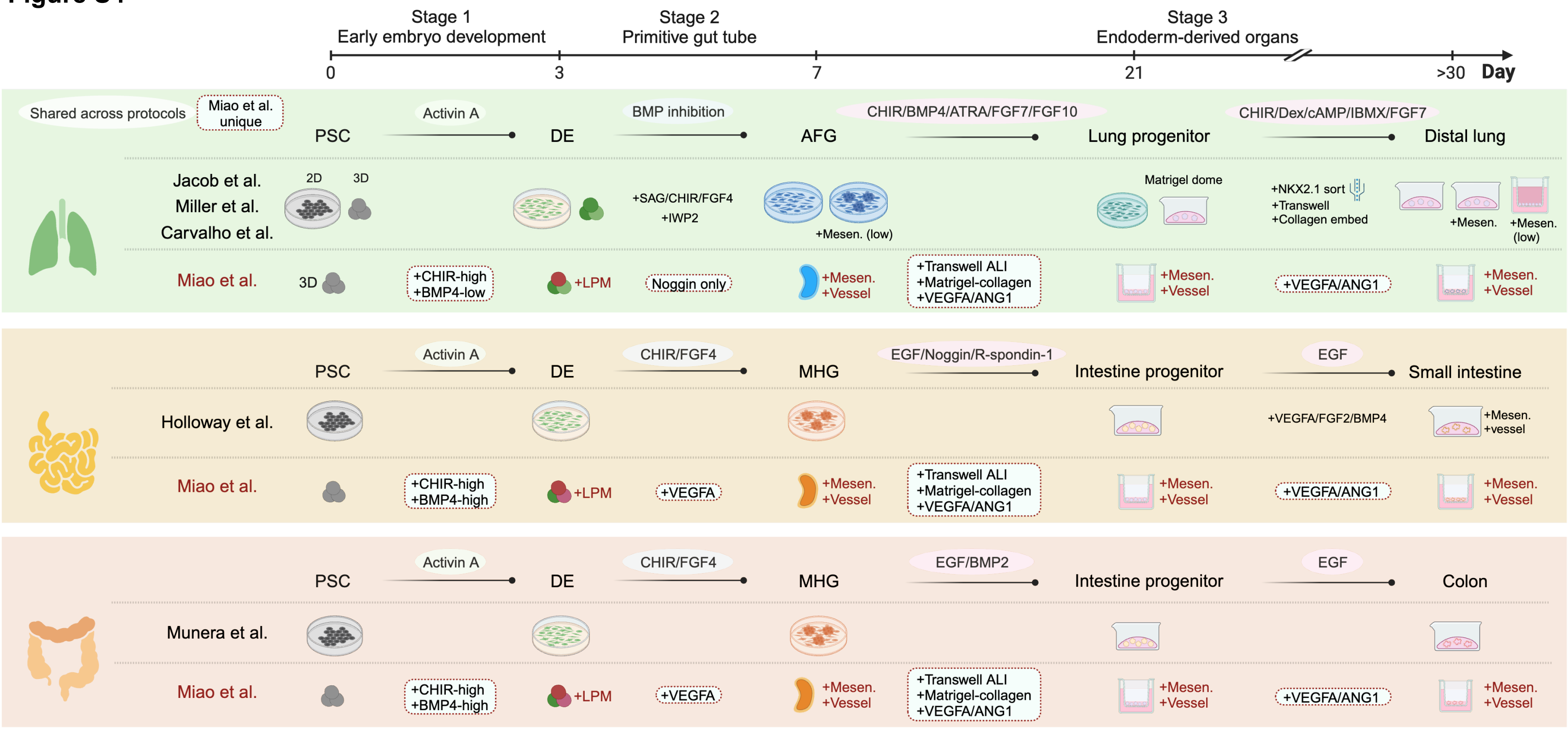


EB: Embryonic body
LPM: Lateral plate mesoderm
DE: Definitive endoderm
AFG: Anterior foregut
MHG: Mid/hindgut
Mesen.: Mesenchyme

CHIR: CHIR99021
ATRA: All-trans retinoic acid
Dex: Dexamethasone
ALI: Air-liquid interface

#### Figure S5

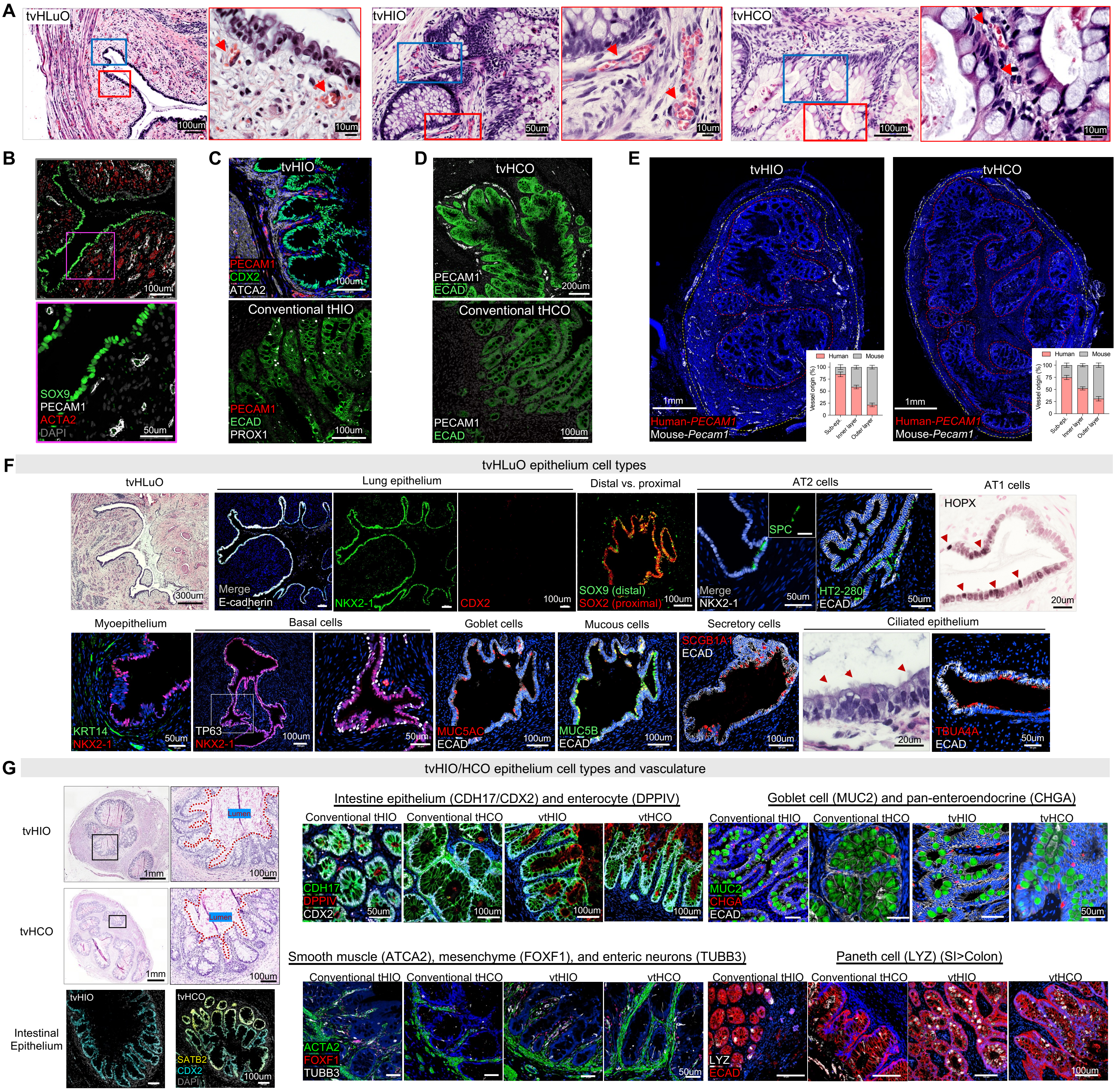

A
tvHLuO
100um
10um
tvHIO
50um
10um
tvHCO
100um
10um
B
100um
SOX9
PECAM1
ACTA2
DAPI
50um
C
tvHIO
PECAM1
CDX2
ATCA2
100um
Conventional tHIO
PECAM1
ECAD
PROX1
100um
D
tvHCO
PECAM1
ECAD
200um
Conventional tHCO
PECAM1
ECAD
100um
E
tvHIO
1mm
Human-PECAM1
Mouse-Pecam1
Human
Mouse
Vessel origin (%)
Sub-epi.
Inner layer
Outer layer
tvHCO
1mm
Human-PECAM1
Mouse-Pecam1
F
tvHLuO epithelium cell types
tvHLuO
300um
Lung epithelium
Merge
E-cadherin
NKX2-1
CDX2
100um
Distal vs. proximal
SOX9 (distal)
SOX2 (proximal)
100um
AT2 cells
SPC
Merge
NKX2-1
50um
HT2-280
ECAD
50um
AT1 cells
HOPX
20um
Myoepithelium
KRT14
NKX2-1
50um
Basal cells
TP63
NKX2-1
100um
50um
Goblet cells
MUC5AC
ECAD
100um
Mucous cells
MUC5B
ECAD
100um
Secretory cells
SCGB1A1
ECAD
100um
Ciliated epithelium
20um
TBUA4A
ECAD
50um
G
tvHIO/HCO epithelium cell types and vasculature
tvHIO
1mm
Lumen
100um
tvHCO
1mm
Lumen
100um
Intestinal Epithelium
tvHIO
tvHCO
SATB2
CDX2
DAPI
100um
Intestine epithelium (CDH17/CDX2) and enterocyte (DPPIV)
Conventional tHIO
Conventional tHCO
vtHIO
vtHCO
CDH17
DPPIV
CDX2
50um
100um
100um
100um
Goblet cell (MUC2) and pan-enteroendocrine (CHGA)
Conventional tHIO
Conventional tHCO
tvHIO
tvHCO
MUC2
CHGA
ECAD
50um
Smooth muscle (ATCA2), mesenchyme (FOXF1), and enteric neurons (TUBB3)
Conventional tHIO
Conventional tHCO
vtHIO
vtHCO
ACTA2
FOXF1
TUBB3
50um
Paneth cell (LYZ) (SI>Colon)
Conventional tHIO
Conventional tHCO
vtHIO
vtHCO
LYZ
ECAD
100um

### Figure S6

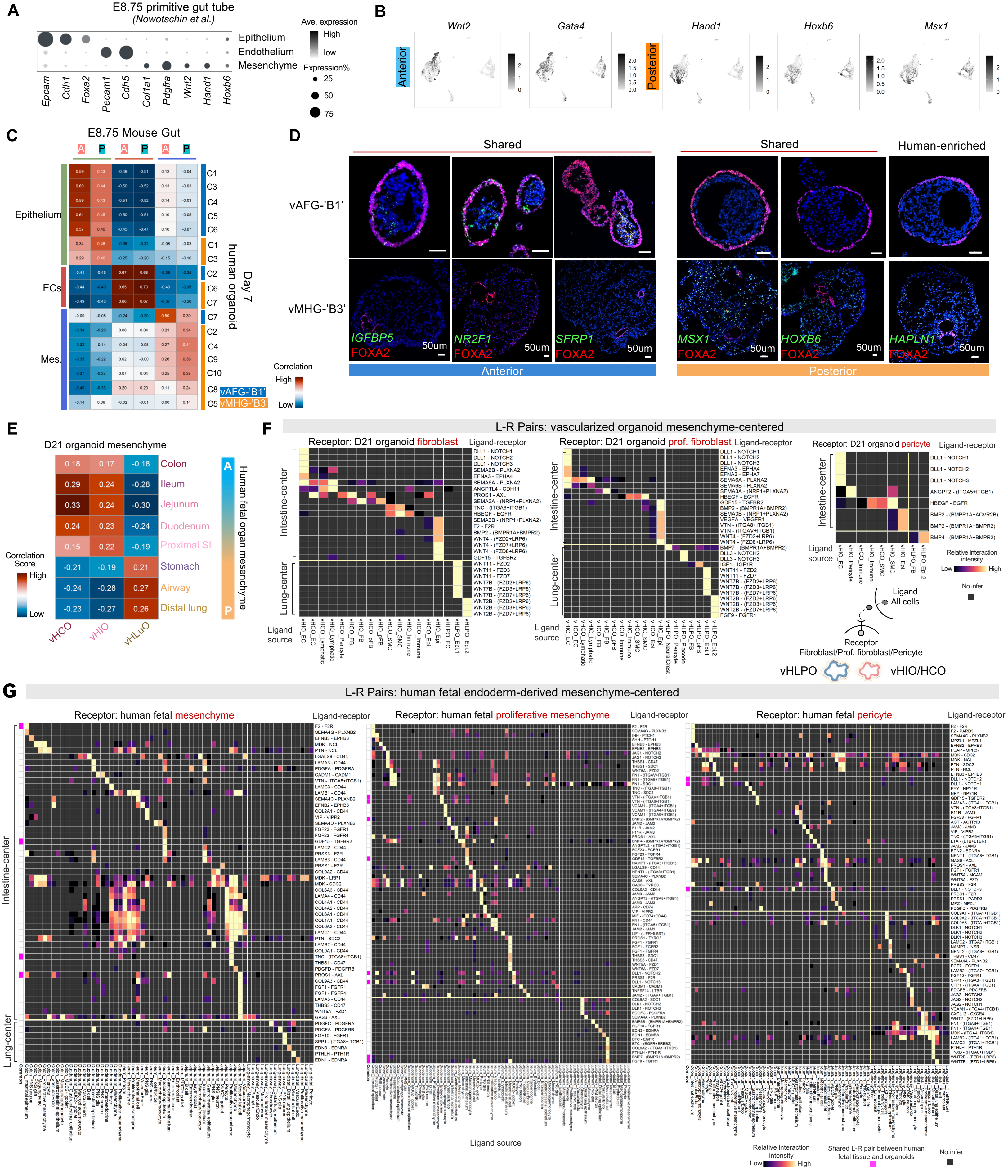

A
E8.75 primitive gut tube
(Nowotschin et al.)
Epithelium
Endothelium
Mesenchyme
Ave. expression
High
low
Expression%
25
50
75
Epcam Cdh1 Foxa2 Pecam1 Cdh5 Col1a1 Pdgfra Wnt2 Hand1 Hoxb6
B
Anterior
Posterior
Wnt2
Gata4
Hand1
Hoxb6
Msx1
C
E8.75 Mouse Gut
Epithelium
ECs
Mes.
human organoid
Day 7
vAFG-'B1'
vMHG-'B3'
Correlation
High
Low
D
Shared
Human-enriched
vAFG-'B1'
vMHG-'B3'
IGFBP5
NR2F1
SFRP1
MSX1
HOXB6
HAPLN1
FOXA2
50um
Anterior
Posterior
E
D21 organoid mesenchyme
Colon
Ileum
Jejunum
Duodenum
Proximal SI
Stomach
Airway
Distal lung
Human fetal organ mesenchyme
Correlation Score
High
Low
vHCO
vHIO
vHLuO
F
L-R Pairs: vascularized organoid mesenchyme-centered
Receptor: D21 organoid fibroblast
Receptor: D21 organoid prof. fibroblast
Receptor: D21 organoid pericyte
Ligand-receptor
Intestine-center
Lung-center
Ligand source
Relative interaction intensity
No infer
Ligand
All cells
Receptor
Fibroblast/Prof. fibroblast/Pericyte
vHLPO
vHIO/HCO
G
L-R Pairs: human fetal endoderm-derived mesenchyme-centered
Receptor: human fetal mesenchyme
Receptor: human fetal proliferative mesenchyme
Receptor: human fetal pericyte
Ligand-receptor
Intestine-center
Lung-center
Ligand source
Relative interaction intensity
Low
High
Shared L-R pair between human fetal tissue and organoids
No infer

### Figure S7

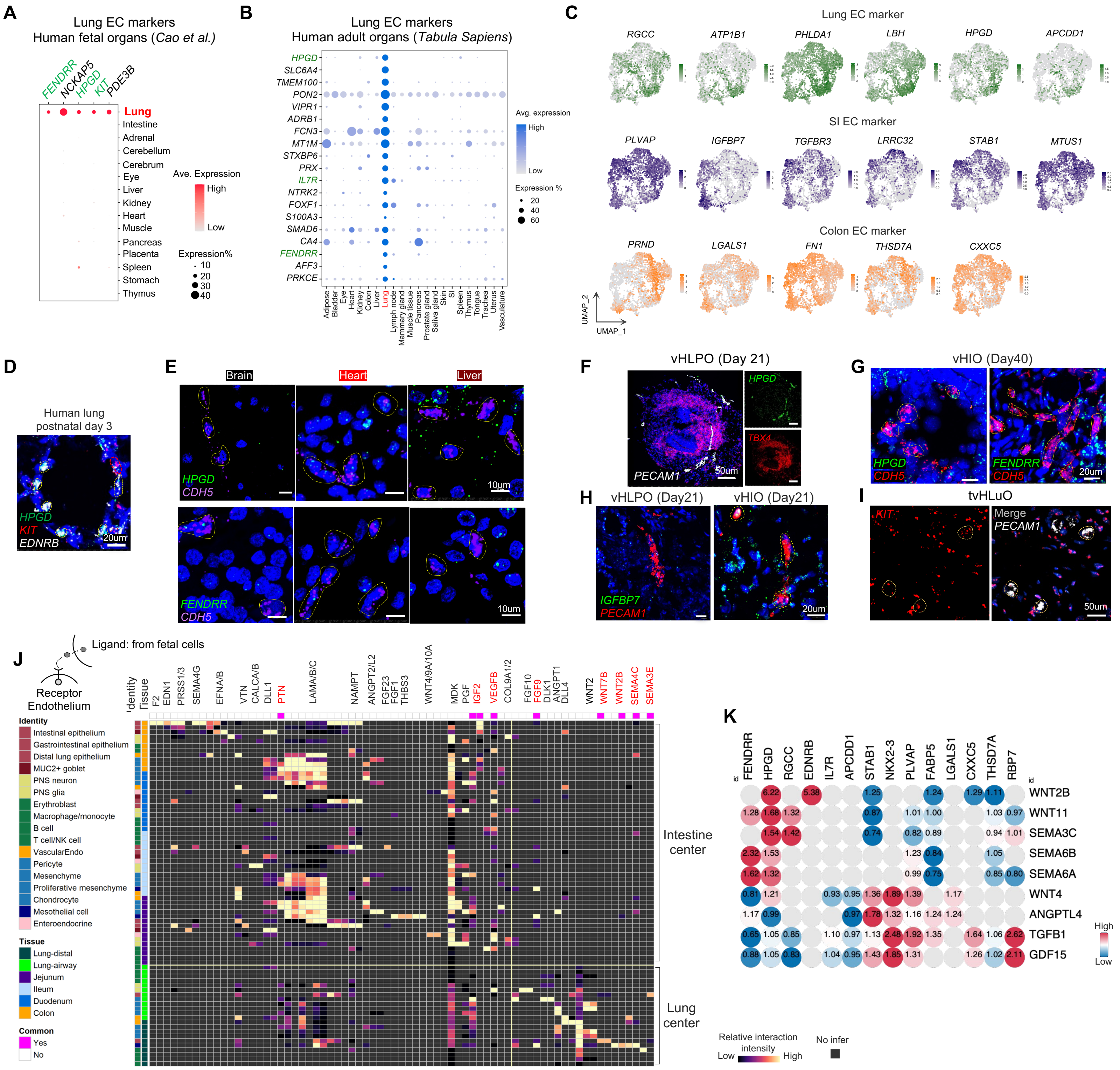
A
Lung EC markers
Human fetal organs (Cao et al.)
FENDRR
NCKAP5
HPGD
KIT
PDE3B
Lung
Intestine
Adrenal
Cerebellum
Cerebrum
Eye
Liver
Kidney
Heart
Muscle
Pancreas
Placenta
Spleen
Stomach
Thymus
Ave. Expression
High
Low
Expression%
10
20
30
40
B
Lung EC markers
Human adult organs (Tabula Sapiens)
HPGD
SLC6A4
TMEM100
PON2
VIPR1
ADRB1
FCN3
MT1M
STXBP6
PRX
IL7R
NTRK2
FOXF1
S100A3
SMAD6
CA4
FENDRR
AFF3
PRKCE
Avg. expression
High
Low
Expression %
20
40
60
Adipose
Bladder
Eye
Heart
Kidney
Colon
Liver
Lung
Lymph node
Mammary gland
Muscle tissue
Pancreas
Prostate gland
Saliva gland
Skin
SI
Spleen
Thymus
Tongue
Trachea
Uterus
Vasculature
C
Lung EC marker
RGCC
ATP1B1
PHLDA1
LBH
HPGD
APCDD1
SI EC marker
PLVAP
IGFBP7
TGFBR3
LRRC32
STAB1
MTUS1
Colon EC marker
PRND
LGALS1
FN1
THSD7A
CXXC5
UMAP_2
UMAP_1
D
Human lung
postnatal day 3
HPGD
KIT
EDNRB
20um
E
Brain
Heart
Liver
HPGD
CDH5
10um
FENDRR
CDH5
10um
F
vHLPO (Day 21)
HPGD
TBX4
PECAM1
50um
G
vHIO (Day40)
HPGD
CDH5
FENDRR
CDH5
20um
H
vHLPO (Day21)
vHIO (Day21)
IGFBP7
PECAM1
20um
I
tvHLuO
KIT
Merge
PECAM1
50um
J
Ligand: from fetal cells
Receptor
Endothelium
Identity
Tissue
F2
EDN1
PRSS1/3
SEMA4G
EFNA/B
VTN
CALCA/B
DLL1
PTN
LAMA/B/C
NAMPT
ANGPT2/L2
FGF23
FGF1
THBS3
WNT4/9A/10A
MDK
PGF
IGF2
VEGFB
COL9A1/2
FGF10
FGF9
DLK1
ANGPT1
DLL4
WNT2
WNT7B
WNT2B
SEMA4C
SEMA3E
Identity
Intestinal epithelium
Gastrointestinal epithelium
Distal lung epithelium
MUC2+ goblet
PNS neuron
PNS glia
Erythroblast
Macrophage/monocyte
B cell
T cell/NK cell
VascularEndo
Pericyte
Mesenchyme
Proliferative mesenchyme
Chondrocyte
Mesothelial cell
Enteroendocrine
Tissue
Lung-distal
Lung-airway
Jejunum
Ileum
Duodenum
Colon
Common
Yes
No
Intestine
center
Lung
center
K
FENDRR
HPGD
RGCC
EDNRB
IL7R
APCDD1
STAB1
NKX2-3
PLVAP
FABP5
LGALS1
CXXC5
THSD7A
RBP7
WNT2B
WNT11
SEMA3C
SEMA6B
SEMA6A
WNT4
ANGPTL4
TGFB1
GDF15
High
Low
Relative interaction
intensity
Low
High
No infer

**Figure S8**

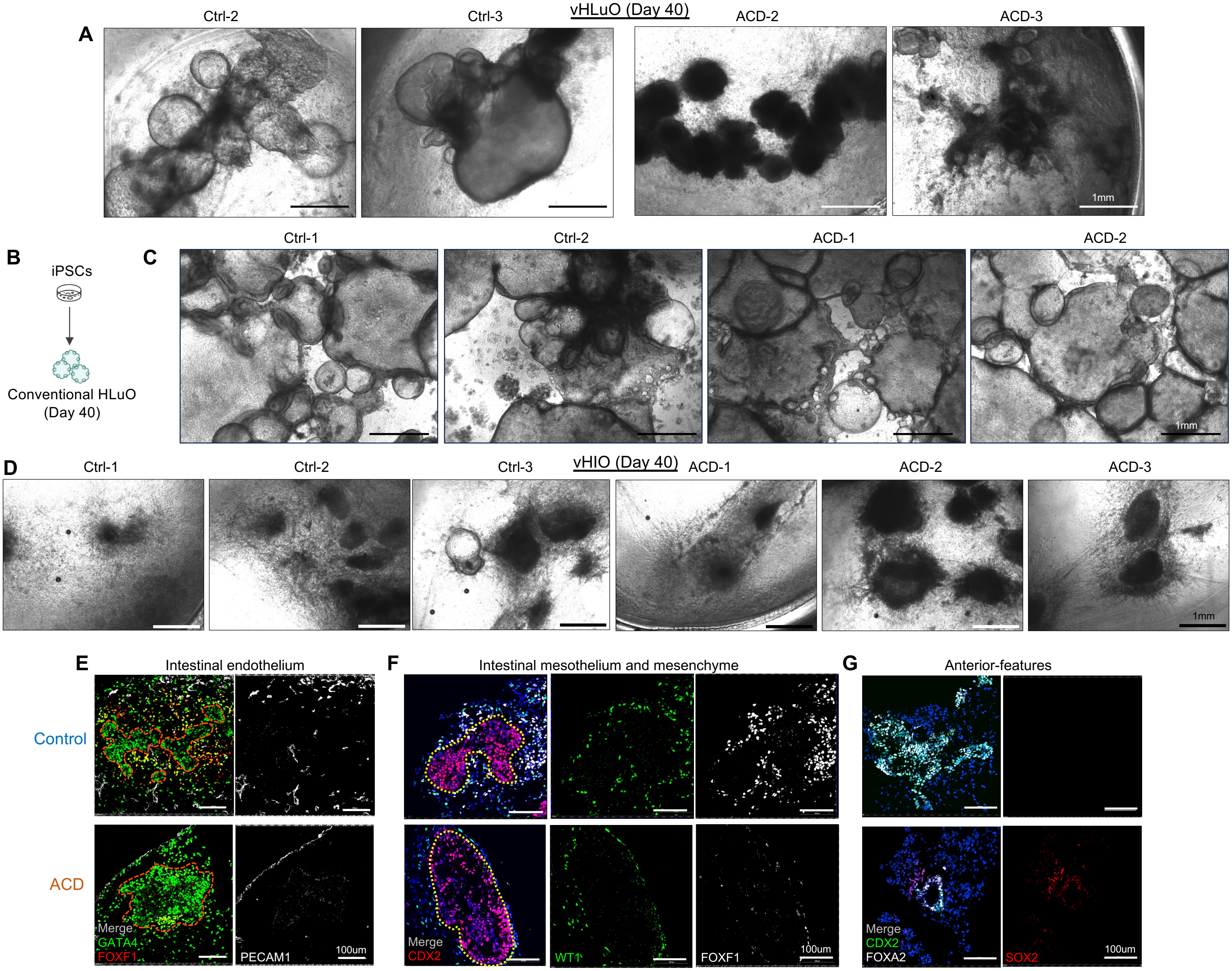

A
vHLuO (Day 40)
Ctrl-2
Ctrl-3
ACD-2
ACD-3
1mm
B
iPSCs
Conventional HLuO (Day 40)
C
Ctrl-1
Ctrl-2
ACD-1
ACD-2
1mm
D
vHIO (Day 40)
Ctrl-1
Ctrl-2
Ctrl-3
ACD-1
ACD-2
ACD-3
1mm
E
Intestinal endothelium
Control
ACD
Merge
GATA4
FOXF1
PECAM1
100um
F
Intestinal mesothelium and mesenchyme
Merge
CDX2
WT1
FOXF1
100um
G
Anterior-features
Merge
CDX2
FOXA2
SOX2
100um
